# A niche-adapted brain microbiome in salmonids at homeostasis

**DOI:** 10.1101/2023.12.07.570641

**Authors:** Amir Mani, Cory Henn, Claire Couch, Sonal Patel, Tomas Korytar, Irene Salinas

**Affiliations:** Center for Evolutionary and Theoretical Immunology, Department of Biology, University of New Mexico, Albuquerque, New Mexico, 87108, USA; Department of Fisheries, Wildlife, and Conservation Sciences, Oregon State University, Corvallis, OR, USA; Norwegian Veterinary Institute, Thormøhlens Gate 53C, 5006, Bergen, Norway; Institute of Parasitology, Biology Centre of the Czech Academy of Sciences, České Budějovice, Czech Republic

## Abstract

Ectotherms have long been known to have peculiar relationships with microorganisms. For instance, bacteria can be recovered from blood and internal organs of healthy teleost fish. However, until now, the presence of a microbial community in the healthy teleost brain has not been proposed. Here we report a living bacterial community in the brain of healthy salmonids. Brain bacterial loads in salmonids are comparable to those found in the spleen and 1000-fold lower than in the gut. Brain bacterial communities share >50% of their diversity with gut and blood bacterial communities. Using culturomics, we obtained 54 bacterial isolates from the brain of healthy rainbow trout. Comparative genomics uncovered unique niche adaptations associated with brain colonization and polyamine biosynthesis. In a natural system, salmonid brain microbiomes shift with the host life cycle, becoming dysbiotic in reproductively mature Chinook salmon, a species that undergoes reproductive death. Our study redefines the relationship between the teleost brain and bacterial microbiomes under physiological conditions. We posit that this symbiosis may endow salmonids with a direct mechanism to sense and respond to environmental microbes.

**One-Sentence Summary:** Salmonids have a brain-adapted, resident bacterial community

## Main Text

Brain-microbiota communication at homeostasis is governed by microbial derived chemical mediators and metabolites that directly or indirectly signal to the brain (*1–3*). During homeostasis, however, viable microorganisms may leak from the gut forming complex microbial communities in gut-distal tissues including lymph nodes and liver (*4–6*). Still, the presence of microbial communities in the vertebrate brain remains highly controversial and only associated with disease states. Evidence for the presence of microorganisms in the diseased human brain is accumulating (*7–10*), but whether brain microbiomes occur at homeostasis remains an unanswered question.

Teleosts appear to be especially permissive to the presence of bacteria in their internal organs during homeostasis. For instance, culturable bacteria could be recovered from the blood and kidney of healthy salmonids (*11*); the biological and functional significance of this observation is still unexplored. More recently, the microbial communities from the spleen of healthy and diseased tilapia were sequenced (*12*), and blood microbiomes have been proposed as a health biomarker in halibut (*13*). This peculiar relationship between teleosts and systemic bacteria is further illustrated by the lack of an endotoxic shock response to lipopolysaccharide injection, a 60-year-old observation that further underscores that teleost internal organs coexist with bacteria (*14*, *15*). Together, these findings motivated us to hypothesize that bacteria and teleosts form symbioses in the brain under physiological states.

Here we show that in salmonids, microbiota directly colonize brain tissues and adapt to this niche. Thus, these findings suggest that microbiota not only regulates the brain via the canonical gut-brain axis but also by directly penetrating the brain. The presence of bacteria in the teleost brain may provide hosts with a direct mechanism for sensing the environment, a finding that redefines how microbes and the brain communicate in the animal kingdom.

## Results and discussion

### Viable bacteria are found in all brain regions of laboratory-reared rainbow trout

Presence of bacteria in the healthy brain is a matter of debate (*16–20*). Under physiological conditions, culturable bacteria can be recovered from the blood and other internal organs of teleosts (*11*, *13*). Thus, we sought to investigate if the teleost brain is also colonized by bacteria at the steady state. We first quantified bacterial levels in four brain regions (olfactory bulb, telencephalon, optic tectum and cerebellum) as well as the gut, spleen and blood of juvenile, laboratory reared rainbow trout. Animals were perfused prior to sampling to remove any blood contamination from the brain. Bacterial loads in the telencephalon (Tel), optic tectum (OT) and cerebellum (Cer) were comparable to those found in the spleen and three orders of magnitude lower than in the gut. In the olfactory bulb (OB), bacterial loads were significantly lower than in the rest of the brain regions examined. In the blood, bacterial levels were the lowest, with 4.6×10^3^ 16S copies/µg tissue or blood DNA compared to 1.9×10^4^ in the spleen (Fig. 1A). Using RNA as a template, estimated bacterial loads were comparable to those using DNA as template (Fig. 1B). The presence of bacterial RNA in the brain suggested that bacteria are viable. To confirm this, we applied culturomics approaches to grow trout gut, blood, spleen, and brain bacterial isolates in different growth media: Luria-Bertani (LB), Nutrient Broth (NB), Tryptic Soy Broth (TSB), MacConkey and Frey Mycoplasma Broth Base; under aerobic and anaerobic conditions, different temperatures (16°C, room temperature, and 30°C) and lysis methods (SI materials and methods). We also obtained cerebrospinal fluid (CSF) from the same animals and plated under the same conditions. Using an NP-40 detergent extraction protocol under either aerobic or anaerobic conditions (Fig. 1C), we recovered 1.9×10^3 to 2×10^3 cfu/g of tissue in the Tel, OT and Cer whereas we obtained 8.9 ×10^2^ cfu/g in the OB. Similar results were obtained when using the mechanical lysis method under aerobic and anaerobic conditions (fig. S1). No bacteria were recovered from CSF samples. A full summary of our culturomics efforts is shown in table S1. Bacterial isolates were identified by 16S amplicon PCR using Sanger sequencing. A total of 54 isolates were recovered from different trout brain regions and different animals (table S1) and 120 isolates were obtained from all tissues sampled. Representative examples of bacterial cultures from the healthy trout brain and other tissues are shown in Fig. 1D. We confirmed that bacteria are localized in the brain parenchyma using fluorescence *in situ* hybridization with universal eubacterial probes (Fig. 1E - H). Combined, our results indicate that the brain of healthy rainbow trout, similar to the blood and spleen, contains viable bacteria at homeostasis.

**Figure 1.**
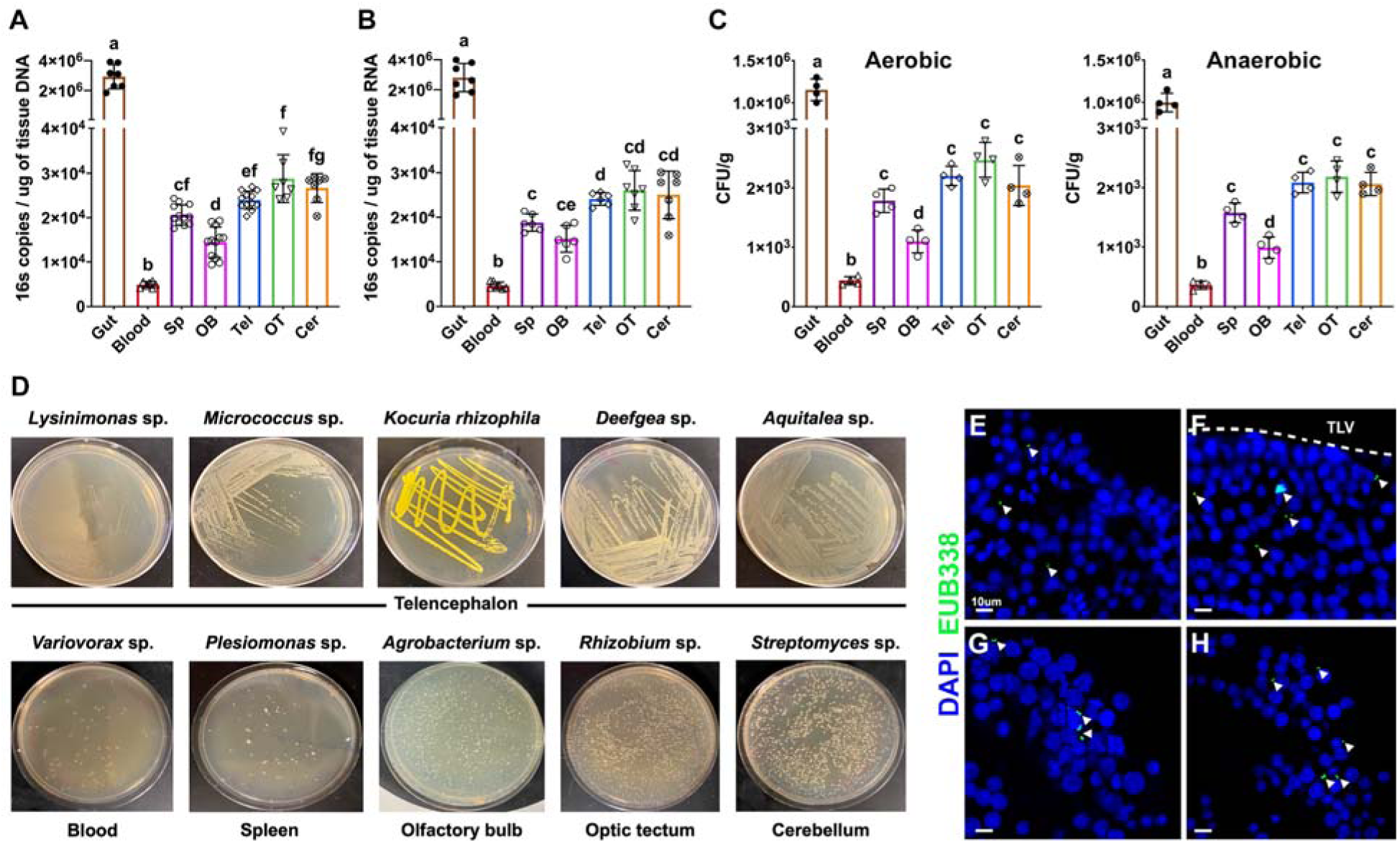
The healthy rainbow trout brain has living bacteria at homeostasis. (A-B) Quantification of 16S rDNA copies in gut, blood, spleen and four brain regions (OB, TL, OT and Cer) of laboratory rainbow trout using DNA (A) or RNA (B) as template. (C) Colony forming units (CFU) per gram of tissue obtained using the NP-40 lysis method under aerobic and anaerobic conditions at room temperature in Tryptic Soy Agar (TSA). Different letters denote statistically significant differences (P<0.05) by Welch’s ANOVA test. (D) Representative examples of different bacterial isolates from control rainbow trout generated via culturomics efforts. (E-H) Fluorescence in situ hybridization of control trout cryosections from different brain regions (E) OB, (F) Tel, (G) OT, and (H) Cer using a universal EUB338 oligoprobe (green). Nuclei were stained with DAPI (blue). Arrowheads indicate bacterial cells. TLV: Telencephalon ventricle.

### Trout brain bacterial communities are only partly sourced by the gut and blood microbiomes

Previous studies in mammals suggest that internal microbiomes originate from leakage of gut bacteria into gut-distant organs (*4–6*). While internal microbiomes in teleosts have been described, their origins are unknown. To resolve this question, we profiled the bacterial communities in all seven tissues sampled in Fig. 1. At the phylum level, the gut bacterial community was largely dominated by Firmicutes, as previously reported in rainbow trout (*21–23*), (Fig 2A-B). In the blood, Proteobacteria and Actinobacteria dominated the community, whereas in the spleen Firmicutes and Proteobacteria were the two dominant phyla. In the four brain regions sampled, Proteobacteria and Actinobacteria were the dominant phyla (63.51%-81.34%) followed by Firmicutes which accounted for 3%-15% of the bacterial community (Fig. 2A). At the family level, Mycoplasmataceae (76.2%) and Chitinobacteriaceae (17.1%) made the majority of the gut bacterial community. Chitinobacteriaceae, however, only represented 1% and 1.8% of the total blood and spleen bacterial communities, respectively, but was detected in all brain samples at similar abundances to the gut. Burkholderiaceae and Propionibacteriaceae, frequently reported in salmonids gut and blood (*24–28*), were only detected in the blood and, at lower abundance, in the brain (Fig. 2B). Enterobacteriaceae accounted for 16.7% and 22.2% of the overall diversity in the blood and spleen, respectively, but were at low relative abundance (<3%) in the rainbow trout brain. PCoA showed tight clustering of the microbial communities from all four regions of the brain and all individuals (Fig. 2C). Water samples clustered apart from all the trout tissue samples sequenced (Fig. 2C, fig. S2). Alpha diversity was lowest in the gut, as previously described in rainbow trout (*21–23*) followed by the blood. The highest alpha diversity was found in the OB and Tel (Fig. 2D and table S2). We next compared the beta diversity of each microbial community to that of the gut and found significant differences in the Weighted UniFrac distances among tissues (Fig. 2E). The greatest distance was observed between the gut and the blood followed by the distance between the gut and each brain region sampled (Fig. 2E). We did not find significant differences when different brain regions were compared to each other. These results suggest that leakage of bacteria from the gut is not the only source of the blood circulating microbiome or the brain microbiome in rainbow trout.

**Figure 2.**
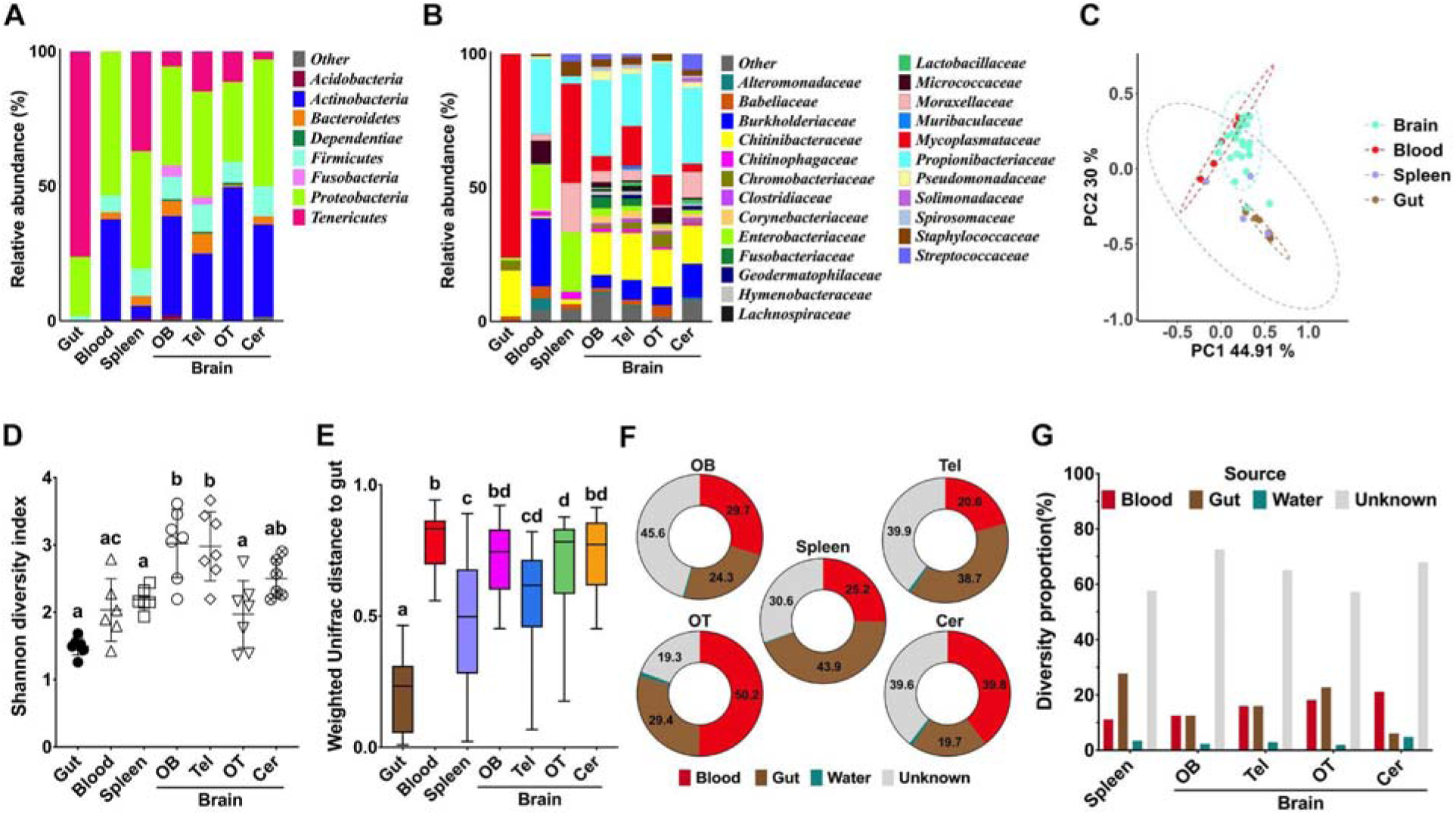
Diversity, composition, and sources of the brain bacterial community in rainbow trout. (A-B) Relative abundance of bacterial phyla (A) and families (B) across the gut, blood, spleen, and four brain regions: olfactory bulb (OB), telencephalon (Tel), optic tectum (OT), and cerebellum (Cer) sampled in this study. (C) Principal coordinate analysis (PCoA) of the rainbow trout gut, blood, spleen, and brain microbial communities. Ellipses represent a 95% confidence interval. (D) Mean Shannon diversity index of rainbow trout gut, blood, spleen, and brain microbial communities. Different letters indicate statistically significant differences (P<0.05) by Kruskal-Wallis test. (E) Weighted UniFrac distance for rainbow trout gut, blood, spleen and the four different brain regions sampled. Different letters indicate statistically significant differences (P<0.05) by Tukey’s post hoc test. (F) Predicted relative percentage of bacterial reads in the spleen, OB, Tel, OT and Cer (sinks) originating from the blood, gut, water or unknown sources using SourceTracker2 analysis. (G) Predicted proportion of the overall spleen and brain (OB, TL, OT and Cer) (sinks) microbial diversity that originates from the blood, gut, water, or unknown sources.

To resolve the source of the brain bacterial community we performed SourceTracker2 analyses using the water, gut or blood as potential sources. Depending on the region, gut sources accounted for 19.7% −38.7% of all reads, whereas the blood accounted for 20.6%-50.2% of all reads in the brain (Fig. 2F-G). Less than 4% of the brain reads originated from the water. Unknown sources not included in our analyses accounted for 20.4%-46% of all reads in the brain, depending on the region. Next, we performed statistical analysis to determine if different brain regions have significantly greater contributions of bacteria of one origin or another. We found that in the Cer, water contributed to the overall diversity at significantly higher proportions than in other brain regions (P-value=0.0137) whereas the greatest unknown sources were found in the OB (table S3, P-value=0.0066). These analyses confirmed that the brain microbial community of rainbow trout can only be partially explained by the gut and blood communities.

In terms of overall diversity, SourceTracker2 predicted that the gut and blood microbiomes combined could only explain up to 40% of the bacterial diversity found in the brain, depending on the region (Fig. 2G). Water contributed to 1.9%-4.9% of the overall brain ASVs diversity. In each of the four brain regions analyzed, we did not detect any differences in the proportion of microbial diversity originating from the gut and blood except for the Cer where the blood’s contribution to microbial diversity was significantly higher (21.2%) compared to the gut (6.1%; P-value= 0.00152, table S3). Combined, our results indicate that the brain bacterial community of healthy rainbow trout is partially sourced from the gut and the blood, with additional unsampled sources or a brain core microbiome accounting for the remaining diversity of the community. These results are supported by culturomics results shown in table S1, with some bacterial species found in the brain but not recovered from the gut, blood or spleen.

### Feeding but not daytime impacts gut and blood bacterial loads but not brain bacterial loads

Previous work in mammals demonstrated circadian oscillation in gut microbiota shaped by feeding-entrained rhythmicity of daily gut IgA secretion (*29*). In teleosts, skin microbiomes display rhythmicity too (*30*). We performed a time series experiment for 30h. Animals were fed at 9 am the first morning of the experiment (fig. S3). We monitored blood glucose levels and observed a peak in blood glucose 17h post-feeding (fig. S3). This slow glucose kinetics are in line with previous reports in trout (*31–33*). Interestingly, bacteria loads associated with the midgut decreased after 12h (fig. S3), the exact same time when food reached the midgut. Decreased gut bacterial loads were paralleled by increased blood bacterial loads at the same time (fig. S3), suggesting perhaps a leakage event from the gut to the blood upon ingesta arrival. In turn, bacterial loads in two regions of the brain examined remained unaltered (fig. S3). These data suggest that feeding and daytime do not affect brain bacterial loads. However, taxa-specific changes were not quantified and therefore we cannot rule out that oscillatory changes in brain microbiome composition occur upon feeding or with circadian cycle.

### Whole genome sequencing (WGS) of brain resident bacterial isolates reveals brain niche adaptations

Pathobionts leaking from the gut into internal compartments undergo within-host evolution in order to adapt to specific niches (*34*). Given the highly different metabolic environments that bacteria are exposed to in the gut, blood and brain, we hypothesized that niche adaptation of microbial isolates of the same species occurs in the salmonid brain. We took advantage of our biorepository of bacterial isolates from rainbow trout gut, blood and brain to perform comparative genomics. We sequenced whole genomes of *Plesiomonas* sp. and *Agrobacterium* sp., both well-known members of the teleost gut microbiome (*35–41*) although *Plesiomonas* sp. has also been reported in diseased teleosts (*42–44*). We sequenced *Plesiomonas* sp. isolates from the gut (n=5), blood (n=5) and brain (n=6) and *Agrobacterium sp*. isolates from the gut (n=5), blood (n=5) and brain (n=4) (table S4 and table S5). All isolates came from two individual fish. We first generated a pan genome for each bacterial species (Fig. 3A and fig. S5A). Interestingly, trout *Plesiomonas* sp. genomes showed astounding diversity depending on their niche. While some isolates (named “long genome” hereafter) contained 4193 genes and 5,020,997 base pairs, others (named “short genome” hereafter) only contained 2615 genes and 3,121,923 base pairs (table S4 and table S5). In the gut and the blood, both short and long genome Plesiomonas isolates were found, whereas in the brain, only long genome Plesiomonas was present (table S4, table S5 and Fig. 3B) suggesting that co-existence of long and short genome Plesiomonas does not occur in the brain niche. Comparative genomic analysis highlighted that while both isolates possess 44% of annotated genes categorized as enzymes, a higher percentage of transporter proteins (12% compared to 6%) and transcriptional regulators (4% compared to 2%) were observed in the long genome *Plesiomonas* sp. (Fig. 3C). This differential gene distribution may confer a selective advantage relevant to the unique environmental pressures within the brain niche. Phylogenetic tree analyses of Plesiomonas genomes from our study and other publicly available revealed that our isolates are more closely related to other fish *Plesiomonas* sp. isolates and that all long genome isolates cluster separately from gut and blood short genome isolates (Fig. 3D). These results are in agreement with core genome and pan genome analyses which identified extensive genetic diversity and the presence of large and variable gene repertoires in *P. shigelloides* (*45*).

**Figure 3.**
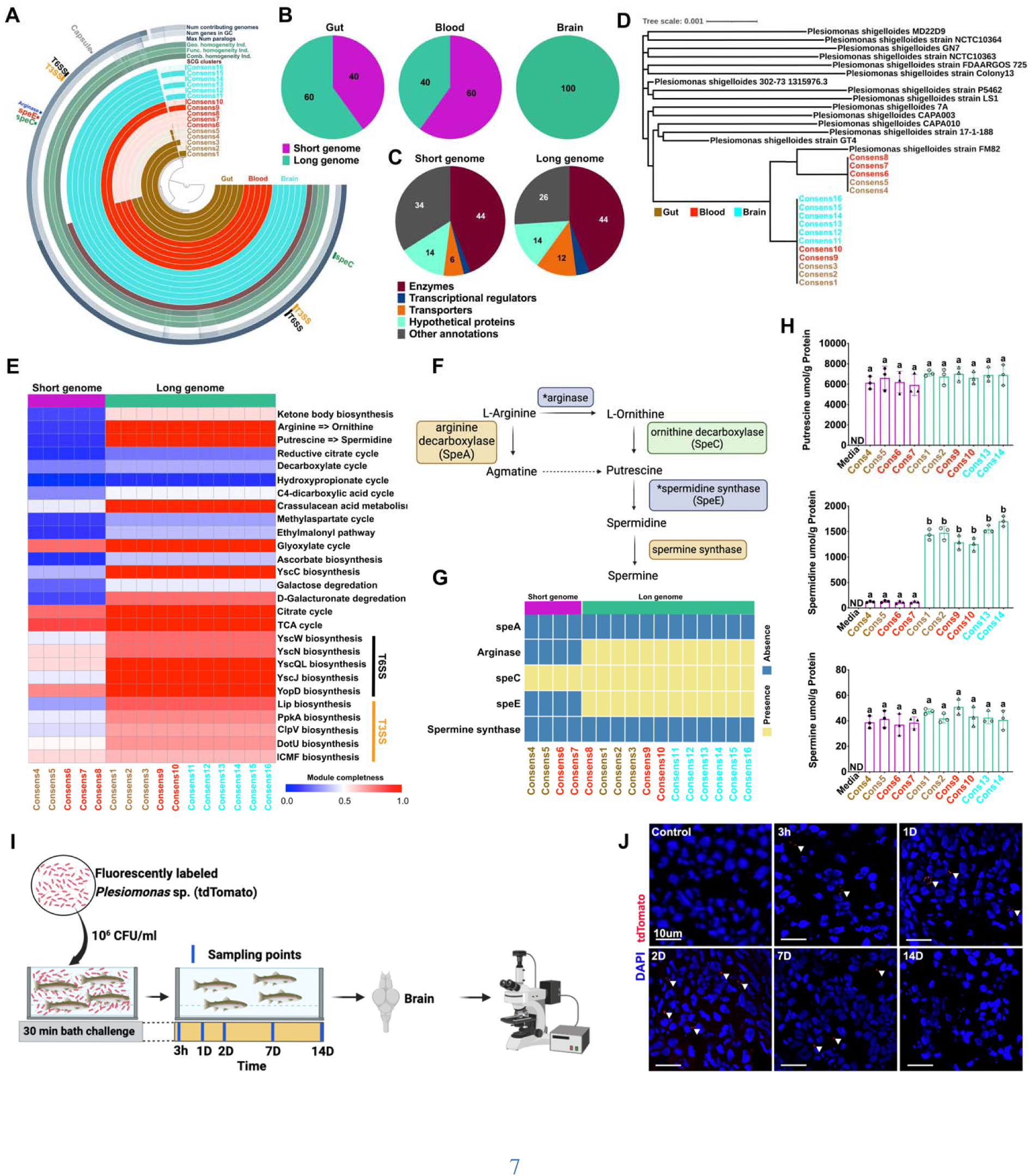
Whole Genome Sequencing (WGS) of brain-resident bacterial isolates reveals brain niche adaptations. (A) Pangenome analysis of sixteen *Plesiomonas*. sp. isolates from gut, blood and brain. (B) Relative percentage of isolates with short and long genomes from each source (gut, blood, brain), highlighting variations in genomic length within species. (C) Functional classes of annotated genes in short genome and long genome *Plesiomonas.* sp. isolates. (D) Phylogenetic tree of *Plesiomonas.* sp. isolates based on whole genome data compared to publicly available strains. (E) Heat map of module completeness for KEGG pathways involved in bacteria polyamines synthesis and secretion systems biosynthesis. (F) Bacterial polyamines synthesis pathway diagram. Asterisks in light purple boxes indicate genes missing in short genome isolates. Green boxes denote enzymes found in all genomes. Light orange box denotes enzymes not found in any of the genomes. (G) Heatmap showing the presence or absence of genes that encode for the main enzymes involved in the polyamine synthesis pathway in each of the Plesiomonas genomes sequenced. (H) Polyamines (putrescine, spermidine and spermine) levels in bacterial growth media (TSB), short genome and long genome Plesiomonas isolates. (ND = Not detectable). (I) Experimental design for bath exposure of juvenile rainbow trout to tdTomato-labeled long genome *Plesiomonas.* sp. gut isolate. (J) Confocal microscopy images of rainbow trout brain cryosections from 3h to 14 days after bath exposure showing presence of tdTomato-Plesiomonas (red, white arrowheads) as early as 3h up until 7 days post-exposure. Images from the OT are shown but bacteria were detected in all brain regions. Cell nuclei were stained with DAPI. Scale bar=20 μm.

Polyamines are produced by both eukaryotes and prokaryotes including the gut microbiota (*46*, *47*). In bacteria, polyamines have essential and non-essential roles in growth, biofilm formation and virulence (*47*). Whereas putrescine and spermidine biosynthesis is widespread among bacteria, spermine production is restricted to some taxa although it can be uptaken from the environment (*48–50*). Microbiota-derived polyamines such as putrescine contribute to host intestinal homeostasis (*46*). KEGG pathway reconstruction of our Plesiomonas genomes identified 39 modules that were significantly enriched in long genome Plesiomonas versus short genome Plesiomonas mostly related to metabolism (Fig. 3E, table S6, fig. S4A). We focused on the stark differences detected in polyamine metabolism modules. All isolates, regardless of the tissue of origin, ubiquitously encode for *odc* (*speC)* (Fig. 3F and G). However, arginase (*arg*), which catalyzes L-ornithine formation from L-Arginine) and *speE* (spermidine synthase, essential for the production of spermidine from putrescine) are only found in long genome isolates (Fig. 3F and G). Nucleotide alignments of those isolates expressing *arg* and *speE* revealed no sequence diversity among brain isolates (file S2). In support, we measured polyamine production (putrescine, spermidine and spermine) by each *Plesiomonas* sp. isolate from ten isolates using ELISA. In order to rule out uptake from the culture medium, we measured polyamine levels in culture medium alone and found them below detection levels in our assay. Our data show that, as predicted from their genomes, all isolates produced similar levels of putrescine and spermine (in low amounts) regardless of the size of their genome and tissue origin, but spermidine production (which is catalyzed by *speE*) was 10-15-fold higher in *Plesiomonas* sp. isolates with long genomes compared to those with short genomes regardless of their tissue origin (Fig. 3H). These results suggest that trout microbiota *de novo* synthesize polyamines and that niche adaptation to the brain may be mediated by acquisition of gene involved in polyamine biosynthesis.

We also identified differences in type 3 and type 6 secretion systems copy numbers, which were more abundant in the long-genome *Plesiomonas* sp. isolates (fig. S4B). Furthermore, the capability to produce capsular components via presence of BDMLBD_07845, GIKMPH_18825 and JOMBDB_16565 genes (Prodigal) (*51*) and to traverse the blood-brain barrier by encoding for endothelial adhesion molecules such as beA, IbeB, IbeC, and IcsA was restricted to the long-genome *Plesiomonas* sp. (fig. S4A). Collectively, our data indicate that adaptations of bacteria to the brain niche consist of: 1) unique abilities to penetrate the blood brain barrier; 2) ability to outcompete other bacteria and/or evade the immune system 3) production of spermidine which, in turn, could regulate cognitive function, neuroprotection, brain immunity and blood barrier permeability (*52–56*). These data underscore the genomic plasticity of *Plesiomonas* sp. within a host, and the ability of this species to adapt to diverse niches with specific nutritional requirements.

In the case of *Agrobacterium* sp. genomes, genome size and number of genes/genome were not different among isolates from different tissues (6016588 base pairs and 5677 genes, fig. S5A-C and table S4 and table S5). Phylogenetic tree analyses indicate that trout-derived Agrobacterium isolates form a distinct clade, diverging as an outgroup relative to the human clinical and plant-based isolates (fig. S5B). Multiple amino acid sequence variation (MAPs) was identified in brain-resident *Agrobacterium* sp. exopolysaccharide biosynthesis gene compared to non-brain isolates (fig. S5E and F). In the gut and blood, all isolates were identical and therefore we called them rainbow trout type strain *Agrobacterium* sp. whereas in the brain, all isolates were named exoP mutants (fig. S5B). exoP (Atu4049, exopolysaccharide polymerization/transport protein) mutants contained 63 amino acid variants but the rest of the genes in this module did not diverge among isolates (fig. S5F). The findings reveal that bacteria capable of infiltrating and persisting in the trout’s brain employ diverse strategies to adapt to this specialized niche.

### Gut-resident bacteria can colonize and persist in the trout brain

Given that 6.2%-22.7% of the diversity of the trout brain bacterial community can be explained from gut sources, we hypothesized that some gut bacterial species could penetrate and colonize the trout brain. We performed an *in vivo* bath exposure experiment in juvenile rainbow trout using a long genome gut *Plesiomonas* sp. isolate fluorescently tagged with tdTomato (Fig. 3I). We detected tdTomato-*Plesiomonas* in all regions of the trout brain by confocal microscopy as early as 3h post exposure, and bacteria persisted at 24h and 48h post-exposure (Fig. 3J). We repeated the same experiment at longer post-exposure times (1 week and 2 weeks) to determine whether colonization occurs long-term. We found long genome tdTomato-*Plesiomonas* in all animals and brain regions after 1 week but bacteria could only be detected at very low numbers after 2 weeks in one of three animals (Fig. 3J). Combined, these results demonstrate short colonization of bacteria in the trout brain by a gut resident *Plesiomonas* sp. isolate. Whether other gut-resident strains are capable of short incursions into the brain as well the functional consequence of this phenomenon remain to be elucidated.

### Geographical survey of brain microbiomes from different salmonid species

In order to extend our observations to other salmonid species and geographical locations, we sampled freshwater and saltwater Atlantic salmon (*Salmo salar*), Gila trout (*O. gilae*), European rainbow trout (*O. mykiss*) and Chinook salmon (*O. tshawytscha*), an anadromous native species of the Pacific Northwest. Sampling locations are shown in Fig. 4A.

**Figure 4.**
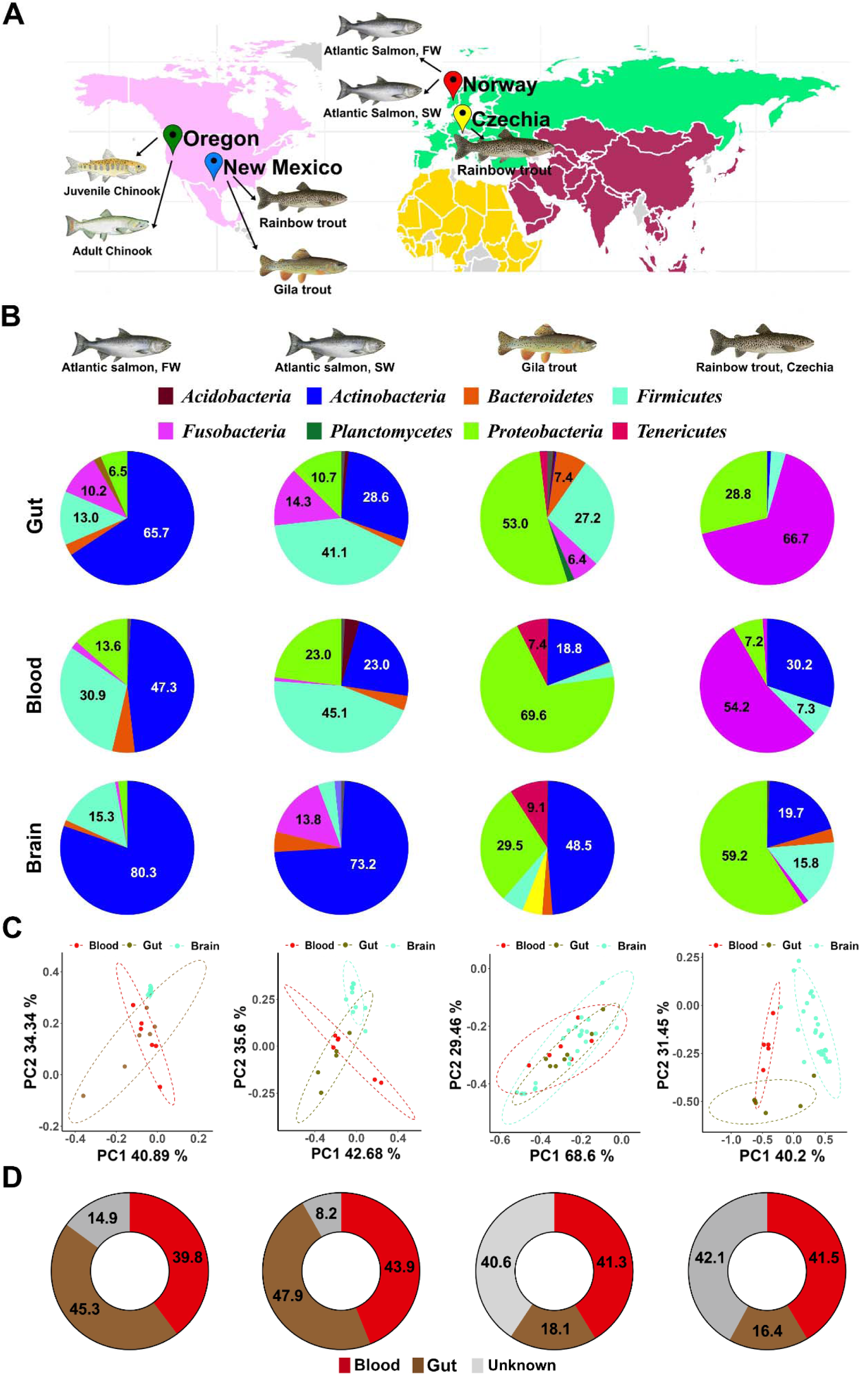
Geographical survey of brain microbiomes from different salmonid species. A) Map of the sampling locations and species sampled in this study. Wild juvenile and adult chinook salmon: Oregon, USA (September 2022). Laboratory-reared rainbow trout and Gila trout: New Mexico, USA. Atlantic salmon (freshwater and saltwater phases). Control laboratory rainbow trout: South Bohemia, Czechia. (B) Relative abundance of bacterial phyla present in the gut, blood, and brain (telencephalon) from freshwater Atlantic salmon, saltwater Atlantic salmon, Gila trout, and rainbow trout from Czechia. (C) Principal coordinates analysis (PCoA) illustrates the microbial community variations within the gut, blood, and brain across in each salmonid group. Note that for Atlantic salmon, only Tel samples are included. Ellipses represent a 95% confidence interval, underscoring significant community composition differences (P < 0.05). (D) Relative contribution of gut and blood microbial communities as potential sources for bacterial communities in the brain (Tel) microbial community in each of the four salmonid groups predicted by microbial SourceTracker2 analyses.

We sampled Atlantic salmon (*S. salar*) from Norway either from freshwater or saltwater sources. In this sampling, we obtained gut, blood, OB and Tel. In freshwater samples, Actinobacteria was the predominant phylum (65.7%) within the gut microbiome, while in saltwater samples, Firmicutes dominated (41.1%). Intriguingly, in the blood, both Actinobacteria and Firmicutes were present at high relative frequencies, though Actinobacteria was more prominent in freshwater salmon (47.3%) and Firmicutes in saltwater salmon (45.1%) (Fig. 4B). The brain microbiome of Atlantic salmon presented a unique profile, distinct from the gut and blood microbiomes. Actinobacteria was consistently the dominant phylum in the brain, accounting for 80.3% in freshwater and 73.2% in saltwater salmon (Fig. 4B). Community composition at the family level, aloha diversity and Weighted UniFrac distances for all Atlantic salmon samples are shown in fig. S6A-F. SourceTracker2 analysis in freshwater Atlantic salmon predicted that 45.3% of ASVs present in the brain are of gut origin, while 39.8% are attributable to the blood (Fig. 4D). In saltwater salmon, gut and blood sources were estimated to contribute 47.9% and 43.9% of the brain’s bacterial ASVs, respectively (Fig. 4D).

Gila trout is a native endangered salmonid species of New Mexico that has undergone severe bottlenecks and genetic drifts through ecological challenges and habitat deterioration (*57–59*). At the phylum level the Gila trout’s gut microbial community was predominantly dominated by Proteobacteria (53%) and Firmicutes (27.2%) (Fig. 4B). At the family level Lactobacillaceae, particularly *Lactobacillus salivarius*, and Chromobacteriaceae were the primary constituents of the gut bacterial community (fig. S6G). In the blood and spleen, Proteobacteria families like Chromobacteriaceae and Betaproteobacterialis were the most frequent taxa. Remarkably, we detected a significant proportion of Propionibacteria constituting 16.4% of the overall blood bacterial community (Fig. 4B and fig. S6G). In terms of the brain microbial composition, Actinobacteria from the Burkholderiaceae family and Proteobacteria from the Propionibacteriaceae family were the most abundant members of the Gila trout brain microbiome (Fig. 4B and fig. S6G). Compared to other salmonids, the alpha diversity of the gut community was significantly higher than that of the other tissues sampled (fig. S6H and table S2) and Weighted Unifrac distances were only significantly different between the gut, blood and spleen and the four brain regions (fig. S6I). PCoA analysis showed a pronounced demarcation in microbial community structures between the gut, blood, and brain (Fig. 4C). Notably, this discrete clustering was found in all the salmonid species under investigation except in adult Chinook salmon (Fig. 5C). SourceTracker2 analyses predicted that in Gila trout, 18.1% of the brain bacterial ASVs originate from the gut and 41.3% from the blood, leaving 40.6% to unknown sources (Fig. 4D).

**Figure 5:**
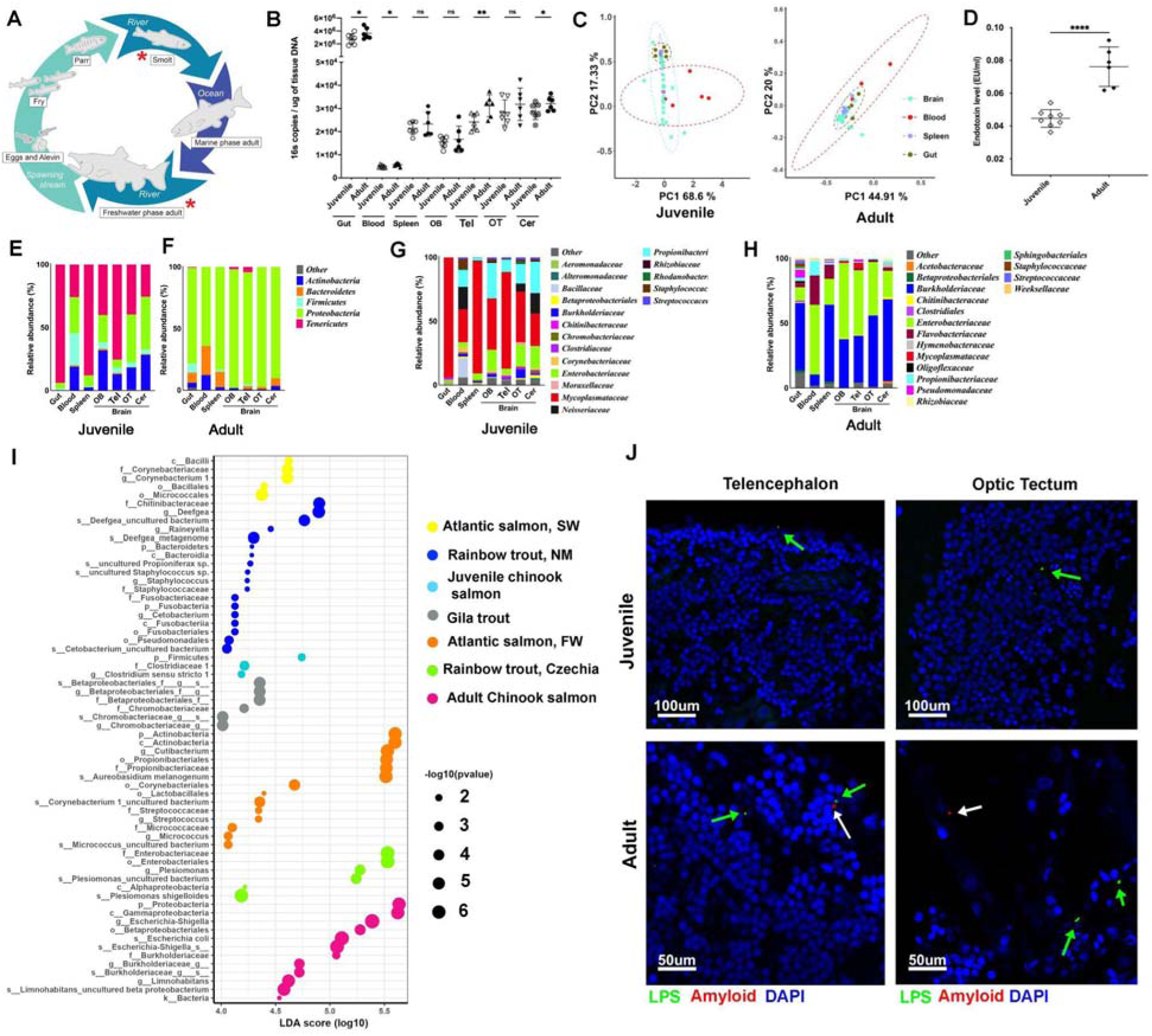
Chinook salmon brain microbiomes shift during life cycle showing signs of dysbiosis in mature individuals. (A) Schematic representation of the natural life cycle of Chinook salmon. Red asterisks indicate the two life stages sampled in this study. (B) Relative bacterial loads quantified by qPCR in the gut, blood, spleen, and four regions of the brain of juvenile and adult Chinook salmon. * P<0.05; **P<0.01 by Welch’s ANOVA test. (C) Principal coordinates analysis of the Weighted Unifrac distance of juvenile and adult Chinook salmon bacterial communities sequenced by 16S rDNA amplicon sequencing. (D) LPS levels in serum from juvenile and adult Chinook salmon, **** P < 0.0001 by Tukey’s post hoc test. (E) Bacterial community composition at the phylum level of the gut, blood, spleen and four areas of the brain of juvenile and adult Chinook salmon. (F) Bacterial community composition at the family level of the gut, blood, spleen and four areas of the brain of juvenile Chinook salmon. (F) Bacterial community composition at the family level of the gut, blood, spleen and four areas of the brain of adult Chinook salmon. (I) Linear discrimination analysis of the brain (Tel only) bacterial communities of all salmonids sampled in this study. (J) Immunofluorescence staining of Tel and OT paraffin sections from juvenile and adult Chinook salmon with anti-*E. coli* LPS antibody (green) and anti-β amyloid (red) shows elevated LPS levels and presence of β amyloid in adult Chinook brain tissues compared to juveniles. Nuclei were stained with DAPI (blue). Green arrows indicate LPS positive puncta and white arrows point at b amyloid positive puncta.

We next sampled European rainbow trout from the Czech Republic, which offered an opportunity to compare their brain microbiome with that of our laboratory rainbow trout from the US. Our results indicate that the bacterial community composition of European rainbow trout was markedly different from the laboratory U.S rainbow trout. This was true for all body sites sampled. Notably, the gut bacterial community of the European specimens contained almost no *Mycoplasma* sp. (fig. S6J). Similar to Atlantic salmon and rainbow trout from the US, the alpha diversity of the gut was significantly lower than that of all brain regions (fig. S6K) and the Weighted Unifrac distance was greatest between the gut, blood and spleen and the four brain regions (fig. 6SL). In turn, the gut community was dominated by Fusobacteria (66.7%) and Proteobacteria (28.8%) (Fig. 4B). In the blood, Fusobacteria (66.7%) and Actinobacteria (30.2%) were the dominant phyla (Fig. 4B). The brain bacterial community was composed of Proteobacteria (32.8%-72.3%, Actinobacteria (9.2%-43.1%) and Firmicutes (15%-22.6%) in different brain regions (Fig. 4B). SourceTracker2 analyses in Czech Republic rainbow trout estimated that 16.4% of the brain bacterial ASVs originates from the gut and 41.5% from the blood, leaving 42.1% to unknown sources (Fig. 4D). A summary of alpha diversity and beta diversity metrics for all microbiome sequences generated in this study is shown in fig. S6 and table S2. Combined, our survey uncovered bacterial communities in the brain of all salmonid species sampled. Salmonid brain microbial communities appear to be largely determined by host genetics as well as environment as is the case for microbial communities associated to mucosal barriers.

### Brain microbiomes shift during life cycle becoming dysbiotic in mature adult Chinook salmon

Chinook salmon are an anadromous salmonid with long, complex, and diverse life histories (*60*). This species has great cultural, economic, and ecological importance in the Pacific Northwest, and many populations are in decline. Understanding the role of the brain microbiome in Chinook salmon is therefore of interest to better understand their physiology and behavior. After spending approximately 8-16 months in their natal freshwater habitats (*61*) juveniles migrate downstream to the ocean where they spend 1-5 years before returning to estuaries and migrating to their natal streams, where they undergo final maturation, reproduce, and die (*60*) (Fig. 5A). The long migration up the river where they will spawn is accompanied by dramatic losses of body condition due to feeding cessation, elevated cortisol levels, lethargy, multi-organ damage, and neurodegeneration (*28*, *62–66*). After spawning, Chinook salmon die in a process known as reproductive death. Previous studies in Atlantic salmon from natural water bodies also captured critical shifts in the gut microbial community during its migratory life cycle (*67*) but shifts in internal microbiomes were not investigated. We hypothesized that adult Chinook salmon microbiomes are markedly different from juveniles and that bacteria may leak from the gut into the systemic compartment and reach the brain, disrupting the healthy brain microbiome. Bacterial loads in the adult gut and blood were significantly higher than in juveniles, whereas the loads in the spleen did not show any differences. In the brain, bacterial loads were significantly higher in telencephalon and cerebellum for sexually mature animals compared to juvenile Chinook salmon (Fig. 5B). Microbiome analyses further supported the convergence of the brain, gut and blood microbial community composition in adult Chinook salmon as shown by PCOA (Fig. 5C). In support, endotoxin levels in serum of reproductively mature Chinook salmon were 3-fold higher compared to juveniles (Fig 5D). As predicted, compared to juveniles, mature Chinook salmon have signatures of pathogenic bacteria circulating in their blood including *Flavobacterium psychrophilum* and *F. columnare* (Fig. 5G-H). Furthermore, we detected greater abundances of Burkholderiaceae and Enterobacteriaceae within the brains of adult Chinook salmon (Fig. 5G-I). Conversely, juvenile Chinook salmon brains were predominantly characterized by the presence of Mycoplasmataceae and Propionibacteriaceae (Fig. 5E-F and 5I). A combined linear discriminant analysis of all the Tel microbial communities for all salmonid species and sampling conditions is shown in Fig. 5I. Since systemic inflammation is known to result in neuroinflammation and breakdown of the blood brain barrier (*68–70*), our findings suggest that the homeostatic brain microbiome becomes disrupted towards the end of the maturation life cycle in Pacific salmon. Whether the changes in microbial signatures detected in adult Chinook salmon brain and blood contribute to the characteristic lethargy, altered swimming behavior (*71*) and neurological dysfunction (*72*) of mature Chinook salmon prior to reproductive death remains unclear.

### Bacteria accumulate in the brain of mature adult chinook next to beta amyloid

Previous work in mammals suggests that amyloid beta plaques are an antimicrobial reaction in the brain (*73–75*). Interestingly, salmonids such as Chinook salmon which undergo reproductive death accumulate beta amyloid in their brain and show signs of neurodegeneration (*66*, *76*, *77*). We therefore hypothesized that beta amyloid deposition in mature salmon may be associated with the penetration of bacteria into the brain. We stained the brains of juvenile and mature adult Chinook with anti-LPS and anti-beta amyloid antibodies. Whereas no amyloid beta and very little LPS signal could be detected in juvenile brains (Fig. 5J), LPS was abundant in the adult Chinook brain, supporting the 16S rDNA sequencing data. Furthermore, LPS was sometimes adjacent to amyloid beta positive regions, especially in the OT (Fig. 5J). This finding suggests a potential association between bacterial dysregulation and amyloid beta accumulation in the brain of mature Chinook salmon, supporting previous findings in mammals.

Microorganisms shape the vertebrate brain via complex biological processes, the best characterized being the gut-brain axis (*1–3*). This bidirectional communication involves molecular mediators released by microorganisms but not direct microbial colonization of the brain. Our findings uncover remarkable associations between the salmonid brain and bacteria during healthy physiological states. Whether this is a hallmark of other teleosts, or a universal symbiotic relationship found in all vertebrates remains to be investigated. Our data demonstrates that members of the salmonid brain bacterial community display niche adaptations both in their genomes and their metabolic capabilities. While our efforts showcase brain niche adaptations of *Plesiomonas* sp. and *Agrobacterium* sp., microbial adaptations to brain niches likely occur in other taxa and span other metabolic functions yet to be discovered. Our work at homeostasis evidently raises the question of how salmonid brain microbial communities are impacted in different disease states and whether they contribute to diverse processes such as sickness behaviors, cognition, locomotion, or sensory perception. In summary, our findings demonstrate that microorganisms can govern the vertebrate brain not only via the gut-brain axis but also by previously undescribed, direct colonization and adaptation to the brain niche; a phenomenon that redefines the boundaries between bacteria and the vertebrate brain.

## Acknowledgments

Authors thank Dr. Darrel Dinwiddie for sharing the Illumina sequencer used in this project University of New Mexico Center for Advanced Research Computer (CARC) for computational resources and data storage support. Authors thanks Dr Sunyer’s laboratory for providing the labeled Plesiomonas strain. Authors thank New Mexico Fish & Wildlife for supplying the Gila trout specimens used in this study, Luke Whitman and other personnel from the Oregon Department of Fisheries and Wildlife (ODFW) for Chinook salmon samples and Dr. Michael Kent for providing the Chinook salmon paraffin blocks. We thank Dr. Marlies Meisel, Dr. Vincent Martinson and Dr. Yi Yang for their helpful feedback on methods and data analysis for this manuscript.

## Author contributions

Conceptualization: IS, AM

Methodology: AM, CC, I

Investigation: AM, CC, CH, IS

Visualization: AM, IS

Bioinformatics analysis: AM

Funding acquisition: IS

Project administration: IS

Supervision: IS

Writing – original draft: AM, IS

Writing – review & editing: AM, IS, CC, CH, SP, TC

## Data and materials availability

All data are available in the main text or the supplementary materials. All 16S ribosomal RNA amplicon sequencing data and bacterial whole genome sequencing have been submitted to the National Center for Biotechnology Information (NCBI) Sequence Read Archive (SRA) under Bioproject accession numbers PRJNA1033688 and PRJNA1032609, respectively.

## Materials and Methods

### Animals

While all animals were monitored for health and appeared healthy during the study, it must be noted that adult Chinook salmon often exhibit opportunistic infections towards the end of their life cycle. This is likely due to increased stress and immunosuppression commonly observed in these fish as they approach reproductive death (*62*, *78*). Laboratory rainbow trout were obtained from Trout Lodge (Washington, USA) as 0.5g larvae and maintained at the University of New Mexico Aquatic Animal Facility. Trout were maintained in a recirculating aquarium system at 16°C and under a 12L-12D photoperiod. Trout were fed commercial pelleted diets (Skretting, USA). Animals were sampled at a mean weight of 30g unless otherwise stated. Gila trout (mean weight=10g) were obtained from the New Mexico Fish and Wildlife facility Albuquerque, New Mexico) and maintained at 10-12 C. All rainbow trout and Gila trout procedures were approved by the University of New Mexico Institutional Animal Care and Use Committee under protocol number 22-201239-MC.

Freshwater Atlantic salmon (*Salmo salar*) (mean weight=60 g) were maintained in a recirculating system at 12°C at the Industrial and Aquatic Laboratory (ILAB) facility, Bergen, Norway. Seawater Atlantic salmon (mean weight=150g) were maintained in a flow through system with 34 ppm salinity at 9C at the same facility. Fish were fed with standard feed from Skretting, with 2- and 3-mm sized pellets, for freshwater and saltwater individuals, respectively. Animals were euthanized, bled and perfused as described below.

Hatchery rainbow trout (mean weight=120 g) from the Czech Republic were obtained from a recirculating aquarium system at the University of South Bohemia. Trout were maintained at 16°C under a 12L-12D photoperiod and fed a commercial diet INICIO plus and EFICO Enviro 920 Advance (BioMar, Denmark). Animal procedures were performed in accordance with Czech legislation (section 29 of Act No.246/1992 Coll. on Protection of animals against cruelty, as amended by Act No. 77/2004 Coll.) and approved by the Ministry of Education, Youth and Sport (MSMT-18301/2018-2).

Wild juvenile Chinook salmon were collected in September 2022 by Oregon Department of Fisheries and Wildlife (ODFW) personnel from a screw trap on the North Santiam River near Stayton, Oregon, USA (latitude 44.7960, longitude −122.7792) under the auspices of a NOAA State 4(d) research permit (#26225). Fish were euthanized using 200ppm tricaine methanesulphonate (MS-222) prior to dissection and tissue sampling. Fresh adult Chinook salmon carcasses were obtained from an ODFW hatchery and sampled on site. These fish were trapped in June of 2022 during their migration up the Willamette River, treated prophylactically with oxytetracycline to prevent bacterial diseases, and transported to holding ponds at Willamette Hatchery near Oakridge, Oregon, USA (latitude 43.9249, longitude −122.8078) for use as brood fish. Adults were held until mid-September, when they were humanely euthanized and artificially spawned for propagation by ODFW employees. Blood and tissues were sampled immediately after euthanasia. All Chinook salmon sampling was approved by Oregon State University Institutional Animal Care and Use Committee (ACUP #2020-0119).

### Tissue sampling and aseptic techniques

Rigorous procedures were implemented to eliminate the possibility of any microbial contamination in brain samples. Animals (n=7 per species) were euthanized with MS-222 (200 ppm, Syndel), and aseptic techniques were strictly followed to collect all samples. Prior to sampling, all incision sites or needle entry points were disinfected using bleach, followed by ethanol cleansing. To ensure a sterile environment, all sampling was conducted adjacent to a Bunsen burner flame. Following blood collection from the caudal vein, a thorough perfusion with 10 ml of sterile PBS was conducted to clear any remaining blood from the tissues. To guarantee no recirculation towards the brain, branchial arch veins and the caudal vein were severed prior to perfusion, enabling a unidirectional flow from the heart to the brain. Autoclaved instruments were employed to collect each tissue sample from the midgut, blood, spleen, and specific brain regions: olfactory bulb (OB), telencephalon (Tel), optic tectum (OT), and cerebellum (Cer). All collected samples were immediately placed in 1ml of sucrose lysis buffer and preserved at −80°C for subsequent processing. Negative controls were included at every stage of the procedure as recommended for low biomass microbiome sequencing (*17*, *79*). Specifically, we included three control tubes containing SLB which were left with their caps open on the same rack as the experimental samples during the sampling procedure. These negative controls were included in every sampling and processed for DNA extraction in the same way as all experimental samples. During transport from Europe to the US, temperature was monitored to ensure that all samples stayed at −80°C.

Animals were euthanized in MS-222 (200 ppm, Syndel). Prior to any sample collection, all sites of incision or needle insertion on the fish were thoroughly disinfected with bleach and subsequently underwent two rounds of ethanol cleansing. All the sampling steps were performed by a Bunsen burner flame. Following caudal vein blood collection, hearts were perfused with 10 ml of sterile PBS to remove any blood contamination in the sampled tissues. The midgut, blood, spleen, and four distinct brain regions: olfactory bulb (OB), telencephalon (Tel), optic tectum (OT), and cerebellum (Cer) were collected and placed in 1ml of sucrose lysis buffer and stored at −80°C until the processing time for all species except for Atlantic salmon where only midgut, blood, OB and Tel were sampled. Any remaining fecal contents were removed during the dissection process prior to collection of the midgut samples.

### Culturomics and colony forming unit quantification

Midgut, blood, spleen, OB, Tel, OT and Cer were collected from a total of eight rainbow trout, which were divided into two groups based on the lysis method applied. Fecal content was removed from the gut before tissue collection. Rigorous aseptic techniques were meticulously adhered to during the sampling process, as explained above and as previously described (*80*). Negative controls consisting of plates that were opened and closed next to the flame but without inoculum were also included. For mechanical lysis, tissues were placed in 300μL of sterile PBS containing sterile glass beads and mechanically lysed at a frequency of 20 shakes per seconds for 3 minutes in a TissueLyser II (Qiagen). Negative controls consisted of tubes containing PBS and beads but no sample. The second set of samples were placed in a tissue lysis buffer consisting of 0.05% NP-40 in PBS to which sterile glass beads were also added and the tubes were treated as for the mechanical lysis (*80*). Lysates were then centrifuged at 1000 rpm for one minute and 30μL of the lysate supernatant were used for culturing under different conditions. We tested Luria-Bertani (LB), Nutrient Broth (NB), Tryptic Soy Broth (TSB), MacConkey and Mycoplasma growth media. All expansion cultures, in conjunction with the plates corresponding to each specimen, were consistently maintained at two distinct temperatures: 16°C, ambient room temperature (25°C) and 30°C. Unique bacterial colonies exhibiting distinct morphologies were meticulously selected from each plate. These colonies were then subjected to 16s rDNA sequencing for bacterial identification, as explained in Phelps et al (*80*).

### DNA extraction and 16S rDNA library preparation

Initial lysis was achieved using two tungsten carbide beads for each tube and subsequently agitated using the Qiagen TissueLyser II for 5 minutes at a frequency of 30 shakes per second. Following mechanical lysis, 200 μl of 1% CTAB (hexadecyltrimethylammonium bromide, Sigma) and 3 μl of proteinase K (100 mg/ml) were added to each sample. DNA extractions were performed as described before Mitchell & Takacs-Vesbach (*81*). The integrity and concentration of the DNA for each sample was measured in a Nanodrop ND 1000 spectrophotometer. Depending on primary concentration, DNA samples underwent dilutions of either 1:10 or 1:20 to use as templates.

Negative and positive controls were included in all our library preparations as recommended by the microbiome research community (*17*, *79*). The DNA template for each PCR reaction was standardized to a minimum of 200 ng to reduce the impact of potential contaminants. Although negative controls did not reach this concentration the product from the DNA extraction step was subjected PCR amplification. Each sample was run in triplicate PCR reactions and the products were pooled prior to cleaning and library preparation. In each library run, a positive control containing DNA from seven known bacterial species grown in the laboratory was included to benchmark sequencing fidelity. An additional control consisting of a mouse colon sample was included in each run to assess consistency across the sequencing runs.

For each sample, three separate PCR reactions were carried out on each sample to amplify the V1–V3 regions of the prokaryotic 16S rDNA, using the primers 28F (sequence: 5’-GAGTTTGATCNTGGCTCAG-3’) and 519R (sequence: 5’-GTNTTACNGCGGCKGCTG-3’) as explained before (*21*). Amplification products for each sample were then pooled and purified by AxyPrep Mag PCR Clean-up Kit (Thermo Fisher Scientific). PCR amplicons were then barcoded using the Nextera XT Index Kit v2 Sets A, B, C and D (Illumina). Amplicon concentration across samples was normalized to 200 ng/μl using the Qubit high sensitivity dsDNA assay before pooling. The library was subject to an additional round of purification using the Axygen PCR clean-up kit. Libraries were sequenced in an Illumina NextSeq 2000 platform using the Illumina NextSeq 2000 Reagent Kit (600 cycle) at the Clinical and Translational Sciences Center at the University of New Mexico Health Sciences Center.

### Microbiome data analyses

Sequences were processed using the Quantitative Insights Into Microbial Ecology 2 (Qiime2, v2023.7) pipeline (*82*). The Divisive Amplicon Denoising Algorithm (DADA2) was employed to cluster demultiplexed sequence reads into amplicon sequence variants (ASVs) (*83*). ASVs were subsequently aligned to the Silva 16S rDNA database (v138) for taxonomic classification (*84*). Prior to the core diversity analyses, each sample was standardized to a depth of 3,500 reads which was sufficient to reach rarefaction. A subsequent core diversity analysis factored both temporal and treatment variables. Indices of alpha diversity (Shannon diversity, Chao1, Faith’s PD) and measures of beta diversity (Weighted UniFrac distances) were generated in QIIME2. Beta diversity metric ordination was visualized with PCoA plots, constructed using the Qiime2R package in RStudio version 1.3.959 (*85*). Using the SourceTracker2 plugin within the QIIME2 framework (*86*), we subjected the 16S rDNA sequences from the spleen, OB, Tel, OT and Cer as sinks and the gut, blood and water (in the case of rainbow trout) were used as presumptive sources for the observed ASVs in each cohort. Squeegee (*87*) was used to identify any potential microbial contamination in brain samples.

### Bacterial load quantification

Bacterial loads in each specimen were measured by employing quantitative PCR for the 16S rDNA as described (*88*). The process involved amplifying the bacterial 16S rDNA sequences in triplicate, utilizing primers that were designed to anneal to the V1 to V3 variable regions, as previously described (*21*). The reaction setup for qPCR was composed in a 96-well plate with 2 µl of normalized DNA/cDNA of 10 ng/µl, 2 µl of primers mix, 6 µl of nuclease-free water, and 10 µl of SsoAdvanced Universal SYBR Green supermix (BioRad). The amplification protocol started with an enzyme activation at 94°C for 90 seconds, followed by 33 cycles of denaturation at 94°C for 30 seconds, annealing at 52°C for 30 seconds, extension at 72°C for 90 seconds, and concluded with a final extension at 72°C for 7 minutes and a stabilization at 4°C (*21*) in a Bio-Rad CFX96 C1000 Touch system. The quantification of the 16S rDNA copies in the samples was calculated against a standard curve derived from a serial dilution of *E. coli* 16S rDNA gene copies, ranging from 10^9 to 10 copies for the V1 to V3 regions. Negative controls were included in all PCR plates.

### In situ hybridization and confocal microscopy

Cryo-mounted fish heads were sectioned to 10 µm thickness using the LEICA CM3050-s cryo-sectioning machine. Prior to sectioning, a meticulous decontamination process was executed wherein all brushes and surfaces of the cryo-sectioning equipment underwent a primary treatment with a 10% bleach solution, followed by dual treatments of absolute ethanol. Following sectioning, brain sections were fixed in a 4% paraformaldehyde (PFA) solution for 10 minutes. Following an overnight permeabilization step in 70% ethanol in DEPC water, samples were then hybridized using Cy5-labeled EUB338 (anti-sense variant) or Cy5-labeled NONEUB (negative control) (Eurofins Genomics). Probes (3 µg/ml) were added to a hybridization buffer containing 2 × SSC integrated with 20% formamide (FisherScientific). Hybridization was performed by incubating the slides at 45°C overnight. After washing the slides with probe-free hybridization medium, two other washes in PBS were performed. Nuclei were stained with 1 μg/mL DAPI for 30 min at 37°C (Invitrogen). Slides were mounted in Fluoroshield medium (Sigma-Aldrich) and imaged on a Zeiss LSM 780 confocal microscope with ZEN microscopy software version 3.3.

### Bacterial whole genome sequencing, assembly, annotation and pan genome analyses

To sequence the genomic profiles of the bacterial isolates outlined in table S3, we cultured each isolate using their respective media and temperature conditions, as delineated in the file S1, for 24-hour. High-molecular-weight DNA from the bacterial pellets was extracted with the QIAGEN DNeasy Blood and Tissue Kit. For consistency, we standardized the DNA extracts to a concentration of 50 ng/µl, as measured by the Qubit High Sensitivity dsDNA assay. A 25 µl aliquot of each DNA preparation was sent to Plasmidsaurus (OR, USA) for Oxford Nanopore sequencing. Upon retrieval of the raw FASTq reads, we employed PycoQC version 2.5.2 to ensure quality assurance (*89*). De novo genome assembly for each sample was performed in Shasta version 0.11.1 (*90*). All the sequenced genomes showed a sequencing coverage of at least 40X. Assembled genomic scaffolds exceeding 500 bp were subjected to a rigorous annotation regimen through Prokka (*91*) and FAMSA (*92*). Pan genome analyses was conducted and visualized using Anvi’o, version 7.1 (*93*). Phylogenetic trees were constructed using IQ-Tree (*94*) and KEGG metabolic reconstructions were performed in Anvi’o-7.1 (*93*).

### Bacterial fluorescent tagging and immersion experiments

Plesiomonas bacteria, tagged with tdTomato, were cultured in tryptic soy broth supplemented with 1% NaCl. The specific strain of Plesiomonas was a gut isolate with long genome (table S5, consens2) isolated from laboratory rainbow trout at the University of New Mexico. After labeling, we resequenced the genome to ensure that there was no contamination during the labeling process. Bacteria were grown to an OD=1.0. Rainbow trout (mean weight 3g) were exposed to aquarium water containing 10^6 CFU/ml of tdT *Plesiomonas* sp. for 30 min at 18°C under gentle aeration. Control animals were subject to the same immersion treatment with aquarium water without bacteria. Trout (N=3/time point) were sampled 3 hours, 1 day, 2 days, 1 week, and 2 weeks after immersion. All fish were perfused with sterile PBS to remove residual blood from the vasculature. Immediately after sampling, fish heads were embedded in a cryo-mounting medium and a rapid freezing process was initiated using liquid nitrogen. All solutions used during tissue processing were prepared by nuclease-free water to prevent potential RNA-DNA degradation. Cryosections (10 µm) were fixed for 5 minutes in 4% paraformaldehyde (PFA) followed by two PBS washes each for 3 minutes. Nuclei were stained with DAPI solution (1 μg/mL; Invitrogen) and slices were mounted with Fluoroshield (Sigma-Aldrich).

### Serum LPS levels in juvenile and mature Chinook salmon

Blood serum samples were initially diluted 1:20 in endotoxin-free phosphate-buffered saline (PBS, Millipore). Subsequently, this diluted mixture was incubated at 70°C for 15 minutes. Following the heat treatment, the quantification of endotoxin levels within these samples was conducted employing the Chromogenic Endotoxin Quantification Kit (Pierce), as per manufacturer’s instructions. To ensure the elimination of potential pyrogenic contamination, all consumables employed in the assay, including pipette tips, tubes, and microplates, were certified as nonpyrogenic.

### Bacterial polyamine ELISA assays

Pure cultures of bacterial isolates were incubated in TSB medium for a 24-hour period, after which the bacterial cells were collected by centrifugation at 5500g for 20 minutes. The cell pellet was subsequently lysed using 1% Triton X-100 (v/v) (Sigma) as described (*95*). The polyamine profile of each isolate including putrescine, spermidine, and spermine levels were quantified employing commercially available ELISA kits (MyBioSource) following the manufacturer’s instructions.

### Blood glucose tests

Following the collection of blood samples, the blood glucose was measured immediately. This was carried out using a TrueMetrix self-monitoring blood glucose meter. A single drop of blood obtained from each sample was applied to TrueMetrix blood glucose test strips, which were then read by the meter to give an instant measurement of blood glucose levels.

### Chinook salmon brain immunofluorescence staining

10 µm-thick paraffin sections from juvenile and adult Chinook salmon brains (n=3/group) were deparaffinized and treated with trypsin antigen retrieval solution (Abcam, ab970) at room temperature for 15 minutes. After rinsing 1x PBS for 5 minutes at room temperature, sections were permeabilized in PBT (1x PBS with 0.1% Triton) for 10 minutes at room temperature. Next, the sections were blocked in 1X PBS/0.1% Tween 20 with 1% bovine serum albumin (BSA) for 30 minutes at room temperature with agitation. After blocking, the samples were incubated in Anti-E. coli LPS antibody [2D7/1] (ab35654) at 1:200 and recombinant Alexa Fluor® 647 Anti-beta Amyloid 1-42 antibody [mOC64] (ab300742) at 1:150, both in 1X PBS/0.1% Tween 20 with 1% BSA overnight at 4°C. On day 2, the sections were rinsed three times with 1x PBS for 5 minutes each at room temperature with agitation. Next, the sections were incubated in goat anti-mouse IgG H&L (Alexa Fluor® 488) (ab150113) at 1:300 in 1X PBS/0.1% Tween20 with 1% BSA for 1 hour at room temperature with agitation. The sections were then rinsed three times in 1x PBS for 5 minutes each at room temperature with agitation. Then the sections were incubated in DAPI at 1:500 in tap water for 2 minutes, rinsed and mounted with KPL fluorescent mounting media. Slides were imaged with a Zeiss LSM 780 confocal microscope using the Zen software.

### Statistical analyses

For the quantification of endotoxin levels in the blood serum across various fish groups, data was initially checked for normal distribution using both F-test and Bartlett’s test. Differences between groups were assessed using one-way ANOVA followed by a Tukey’s post hoc test. Bacterial loads, as quantified by 16S gene copies across different tissues, were subjected to the same normality checks. To discern variations among the different tissues and between groups Brown-Forsythe and Welch’s ANOVA test were employed. For the microbiome Shannon Diversity data, normal distribution checks were first conducted as described. Given the non-parametric nature of the data, a Kruskal-Wallis test was applied to determine differences among groups. In case of significant differences, post hoc pairwise comparisons were conducted with appropriate corrections for multiple testing. Weighted UniFrac distances were visually interpreted using PCoA plots. Differences in clustering on these plots across groups were tested for significance using the multivariate dispersion analysis. The data derived from SourceTracker2, representing percentages of various sources, and the module completeness percentages obtained from Anvi’o KEGG Pathways were first subjected to the Shapiro-Wilk Test to assess their distribution for normality. Following this, to investigate the differences between distinct groups, we employed the Mann-Whitney U Test for post hoc analyses. Significance in the presence or absence of specific genes or pathways in the pangenome analysis was evaluated using Fisher’s exact test. All statistical analyses were conducted using the GraphPad Prism (V10.0.3) and RStudio (R 4.2.3). Differences were considered statistically significant when P<0.05.

## Supplementary Text

**figure S1.**
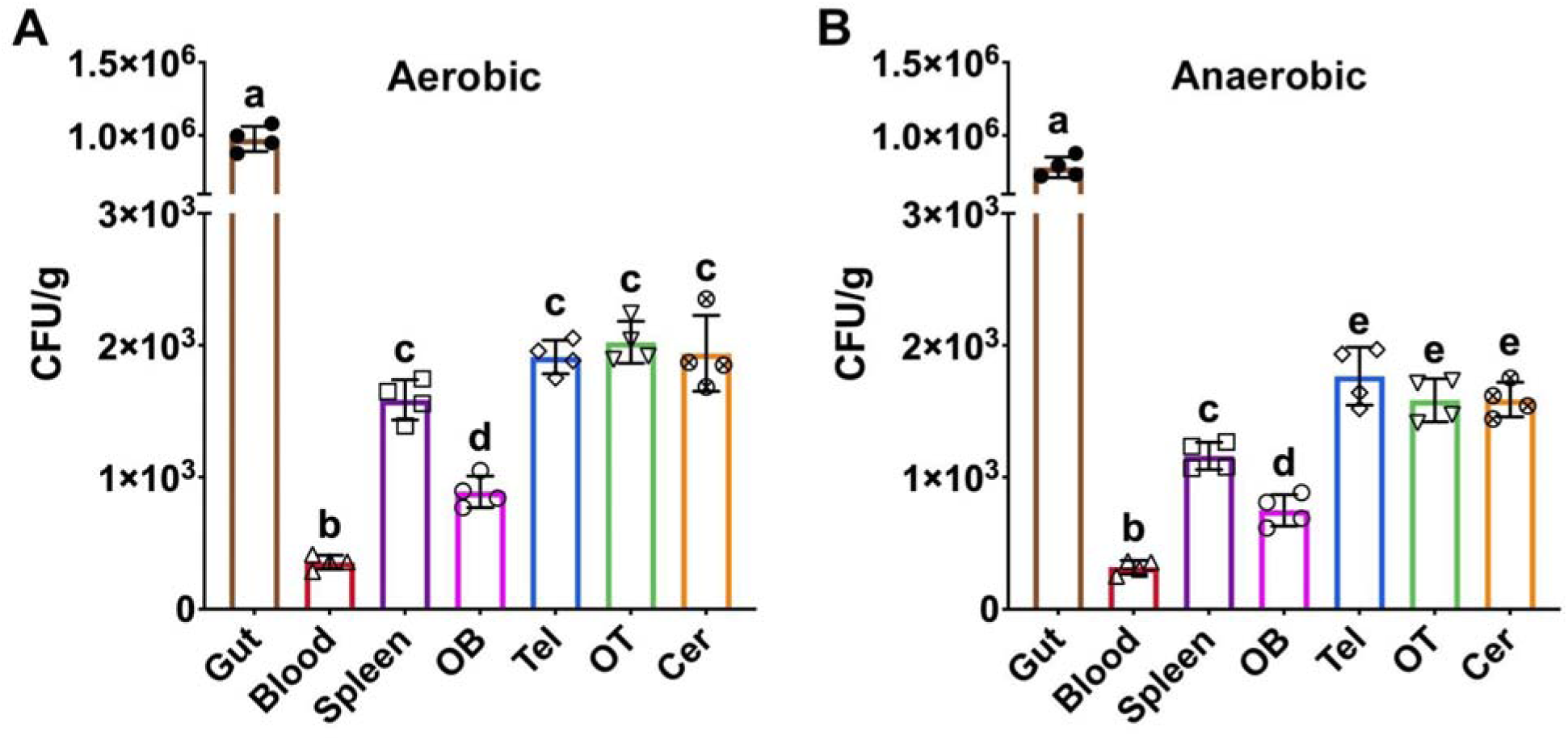
Colony forming units (CFU) per gram of tissue following mechanical lysis from laboratory rainbow trout gut, blood, spleen and brain (OB, Tel, OT and Cer) grown in TSA plates under aerobic (A) and anaerobic (B) conditions. Different letters denote statistically significant differences P<0.05 by Welch’s ANOVA test.

**figure S2.**
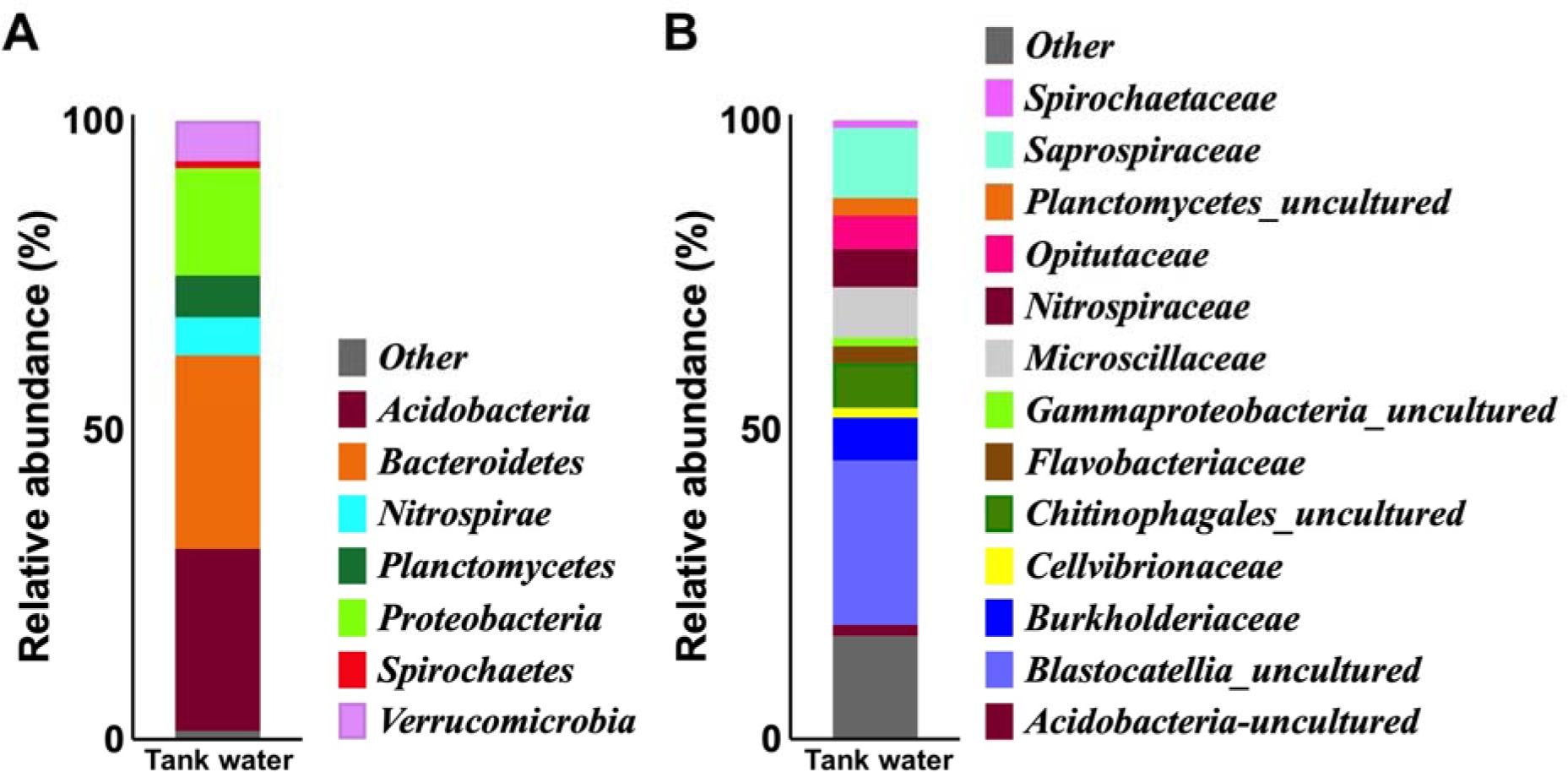
(A-B) Relative abundance of bacterial phyla (A) and families (B) in tank water (n=3) from laboratory rainbow trout.

**figure S3.**
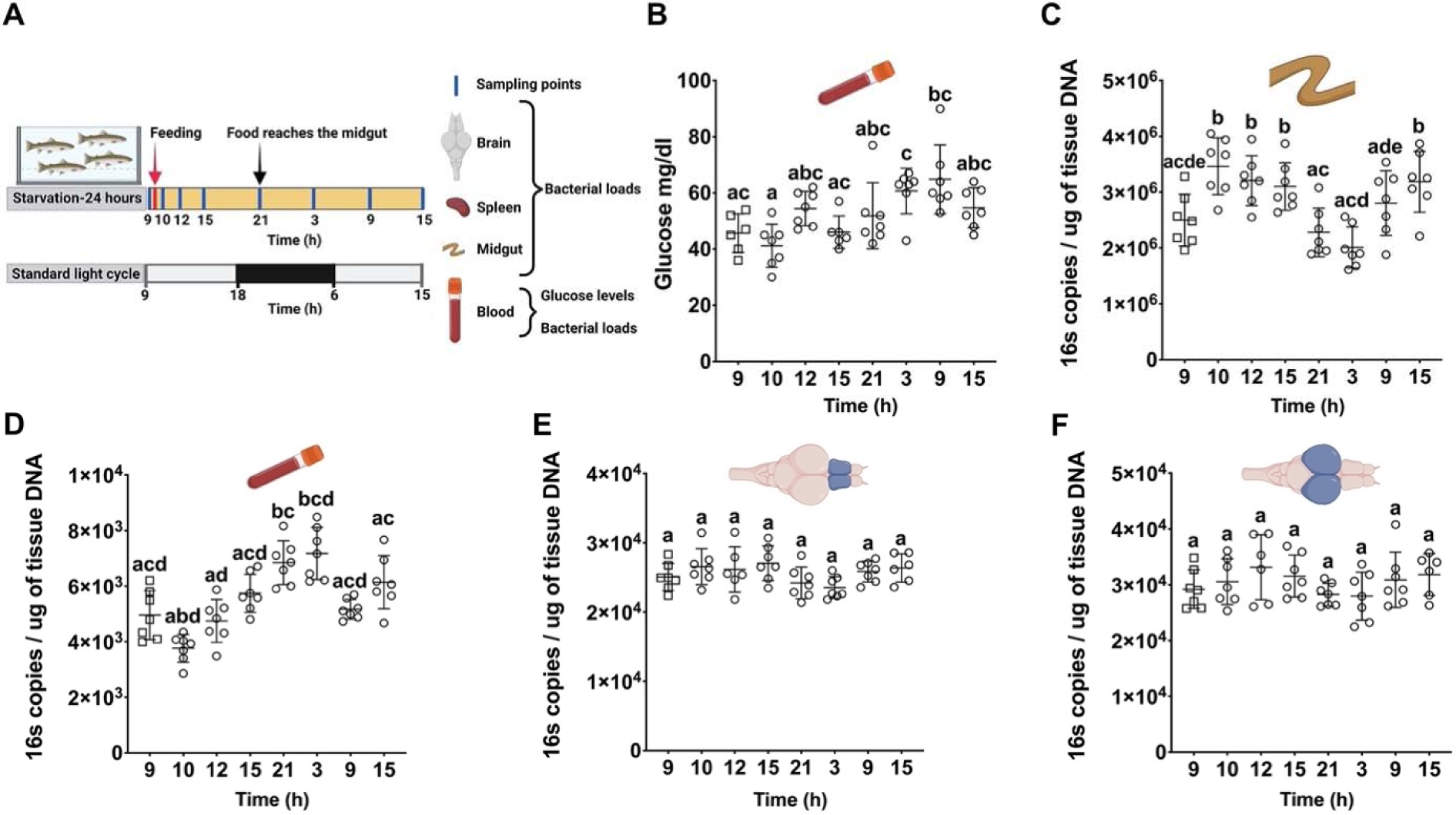
Feeding influences bacterial loads in the gut and blood, but neither feeding nor daytime affect bacterial loads in the brain. (A) Schematic representation of the experimental design and conditions. The x-axis represents the hours of the day, beginning at 9 AM (09:00) on the initial day and concluding at 3 PM (15:00) on the following day. (B). Temporal variation in blood glucose during the time course of the experiment in the fed group. (C-F) Quantification of 16S rDNA copies per microgram of tissue DNA in the gut (C), blood (D), Tel (E) and OT (F) during the course of the experiment. Different letters denote statistically significant difference by Welch’s ANOVA test for 16S loads and (P<0.05) and Tukey’s post hoc test for glucose levels.

**figure S4.**
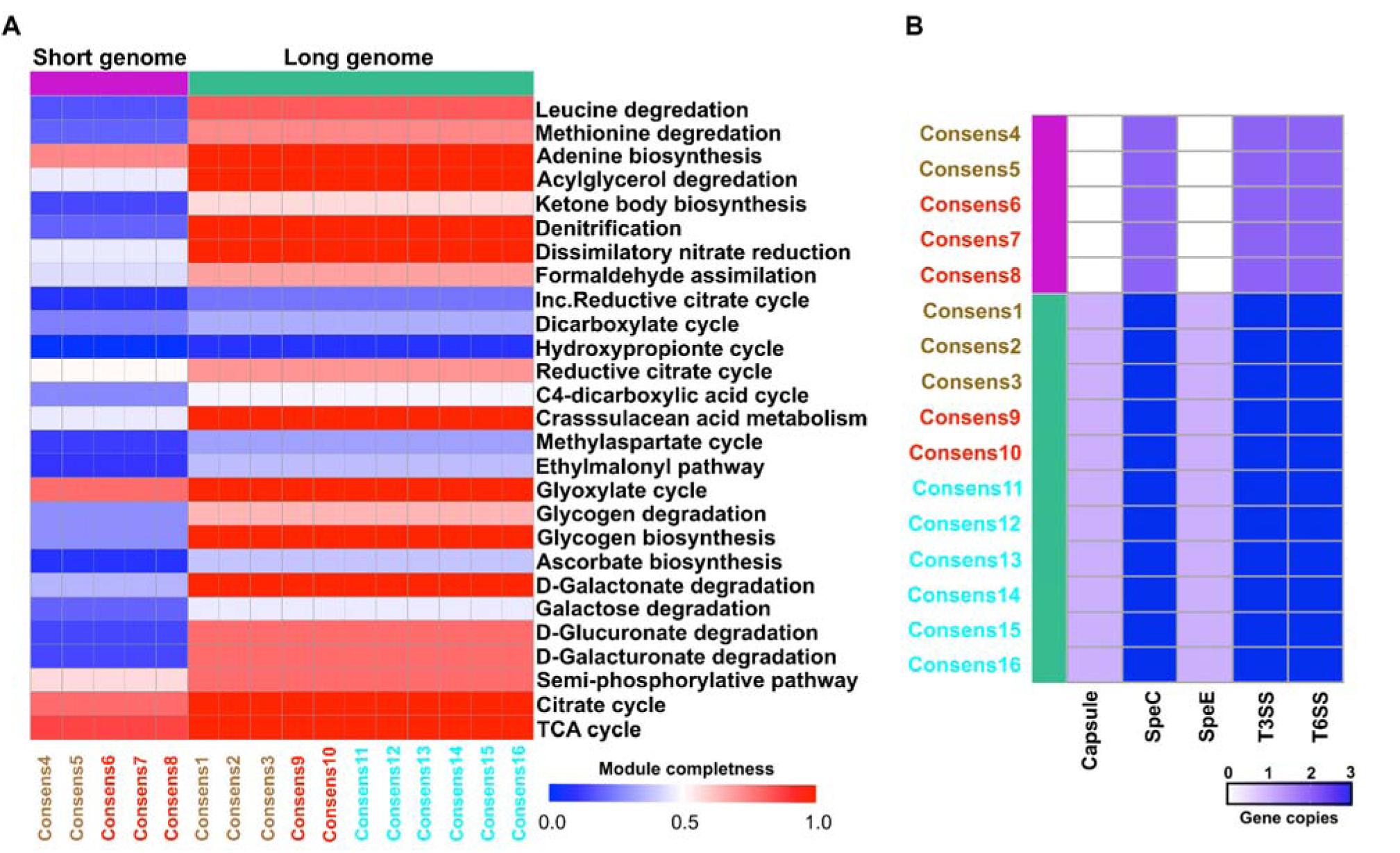
Whole genome sequencing (WGS) of brain-resident bacterial isolates reveals brain niche adaptations. (A) Heat map of top 30 significantly different KEGG pathways between short genome and long genome *Plesiomonas* sp. isolated from various tissues of laboratory rainbow trout. (B) Heat map of gene copy numbers for membrane capsule, polyamines synthesis pathway enzymes and secretion systems for short genome and long genome *Plesiomonas* sp. isolated from various tissues of laboratory rainbow trout.

**figure S5.**
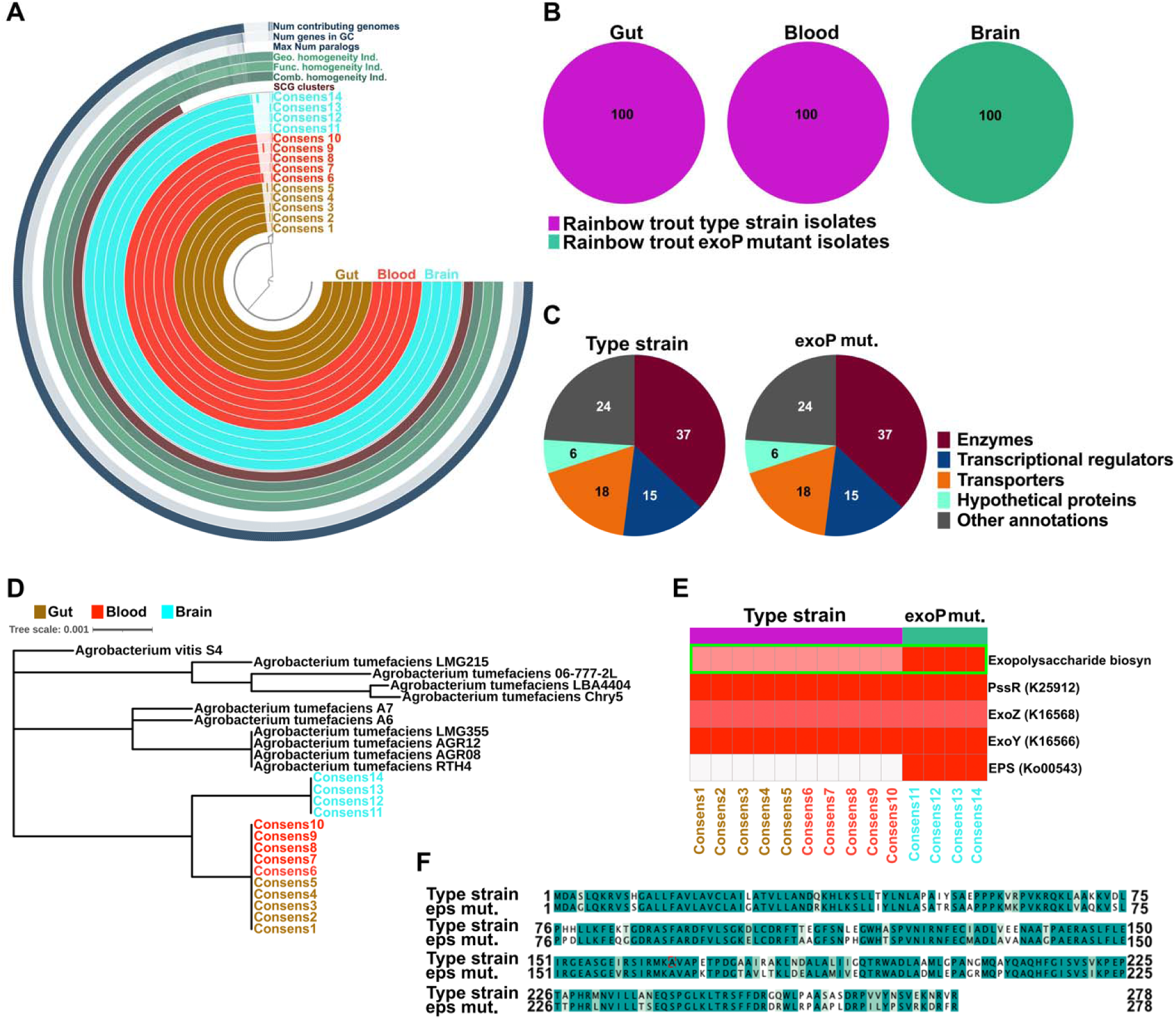
Whole genome sequencing (WGS) of brain-resident *Agrobacterium* sp. reveals brain niche adaptations. (A) Pangenome analysis of fourteen *Agrobacterium* sp. isolates from rainbow trout gut, blood and brain. (B) Relative percentage of rainbow trout type strain and exoP mutant *Agrobacterium* sp. from each source (gut, blood, brain). (C) Functional classes of annotated genes in rainbow trout type strain and exoP mutant *Agrobacterium* sp. isolates. (D) Phylogeny tree of *Agrobacterium* sp. isolates based on whole genome data compared to publicly available strains. (E) Heatmap of module completeness for KEGG indexes involved in *Agrobacterium* sp. exopolysaccharide biosynthesis. (F) Multiple amino acid sequence alignment for rainbow trout type strain and eps mutant *Agrobacterium* sp.

**figure S6.**
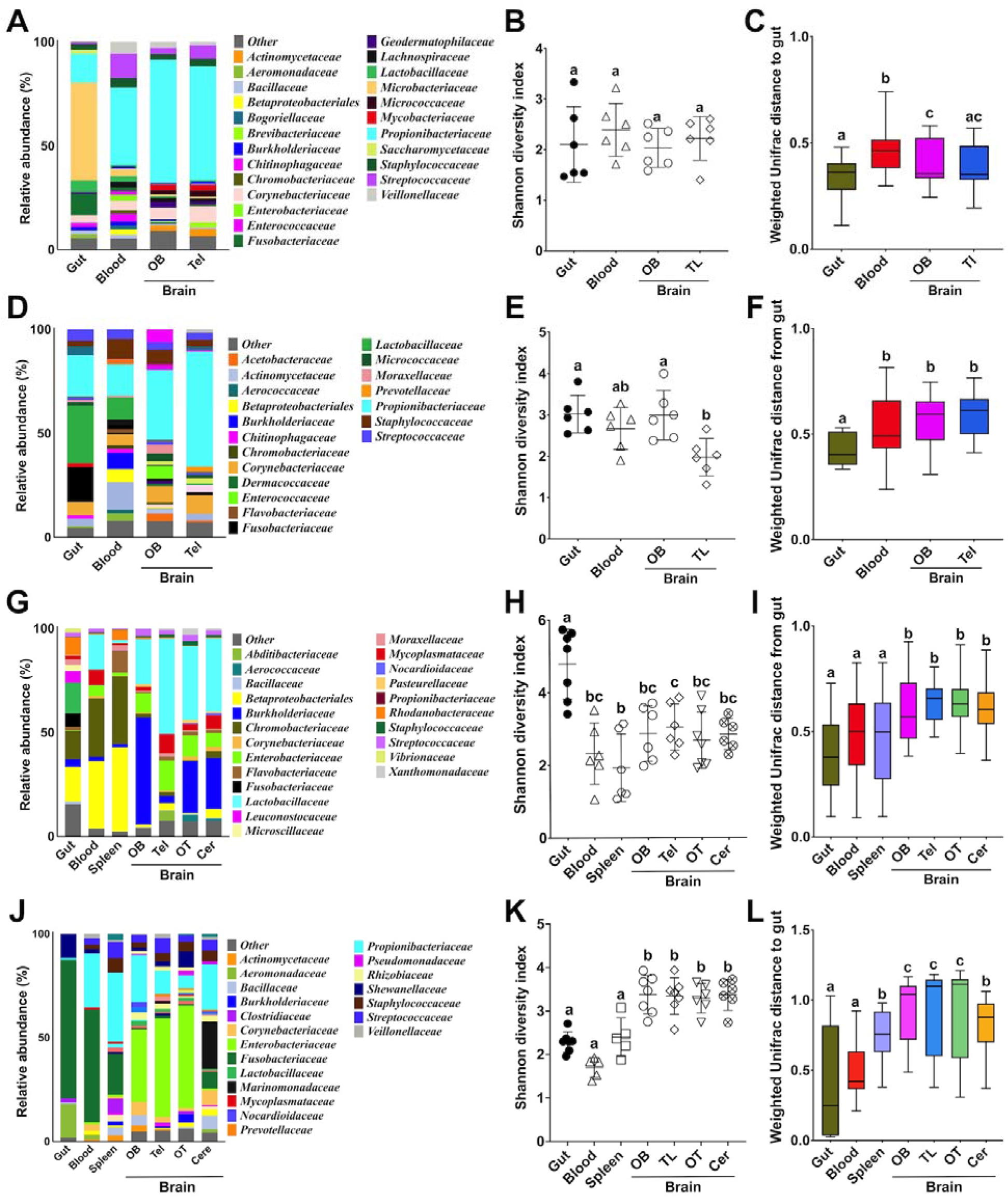
Bacterial community composition analyses of different salmonid species across various tissues. Freshwater Atlantic salmon bacterial community composition at the family level in the gut, blood, OB, and Tel (A). Mean Shannon diversity index for the gut, blood, OB, and Tel of freshwater Atlantic salmon (B). Weighted UniFrac distance for the gut, blood, OB, and Tel of the freshwater Atlantic salmon bacterial communities (C). Saltwater Atlantic salmon bacterial community composition at the family level in the gut, blood, OB, and Tel (D). Mean Shannon diversity index for the gut, blood, OB, and Tel of saltwater Atlantic salmon bacterial communities (E). Weighted UniFrac distances for the gut, blood, OB, and Tel of saltwater Atlantic salmon bacterial communities (F). Gila Trout bacterial community composition at the family level in the gut, blood, and brain (OB, Tel, OT and Cer) (G). Mean Shannon diversity index for the gut, blood, and brain (OB, Tel, OT and Cer) bacterial communities from Gila trout (H). Weighted UniFrac distances for the gut, blood, and brain (OB, Tel, OT and Cer) bacterial communities in Gila trout (I). Bacterial community composition at the family level in the gut, blood, and brain (OB, Tel, OT and Cer) from rainbow trout from Czechia (J). Mean Shannon diversity index for the bacterial communities of the gut, blood, and brain (OB, Tel, OT and Cer) of rainbow trout from Czechia (K). Weighted UniFrac distances of the bacterial communities from the gut, blood, and brain (OB, Tel, OT and Cer) of rainbow trout from Czechia (L). Different letters denote statistically significant groups (P<0.05) by Kruskal-Wallis test.

### Supplementary tables S1-S4

**Table S1.** Tissue distribution of isolated bacteria from laboratory rainbow trout tissues.

■ Np-40 only ■ Mechanical lysis only ■ Np-40 and mechanical lysis
| 16s sequence blast match | Gut | Blood | Spleen | OB | Tel | OT | Cer | Number of isolates |
| --- | --- | --- | --- | --- | --- | --- | --- | --- |
| *Achromobacter insuavis* |  |  |  |  |  |  | □ | 1 |
| *Agrobacterium* spp. | □ | □ | □ | □ | □ | □ | □ | 42 |
| *Aquitalea* sp. | □ |  |  |  | □ |  |  | 2 |
| *Bacillus nealsonii* |  |  | □ |  |  |  |  | 2 |
| *Deefgea* spp. | □ |  | □ |  | □ |  |  | 3 |
| *Escherichia coli* |  | □ |  |  |  |  |  | 1 |
| *Hafnia alvei* |  |  |  |  | □ |  |  | 5 |
| *Kuckoria rhizophila* |  |  |  | □ | □ |  |  | 3 |
| *Lysinimonas* sp. |  |  |  |  | □ |  |  | 1 |
| *Micrococcus* sp. |  |  |  |  | □ |  |  | 2 |
| *Plesiomonas* sp. | □ | □ | □ |  | □ | □ |  | 50 |
| *Rhizobium* sp. |  | □ |  | □ |  | □ |  | 3 |
| *Shewanella* sp. | □ |  |  |  | □ |  |  | 4 |
| *Streptomyces* sp. |  |  |  |  |  |  | □ | 1 |
| *Variovorax* sp. |  | □ |  |  | □ |  | □ | 4 |

**Table S2.**
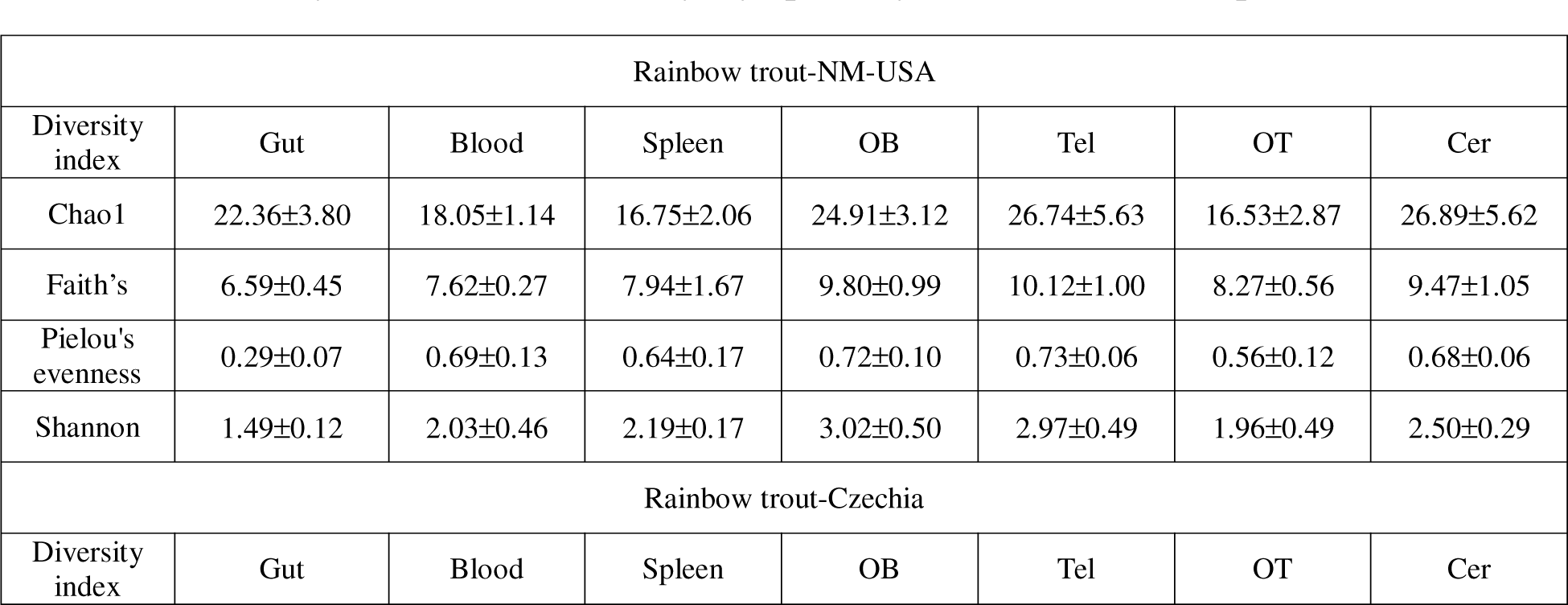
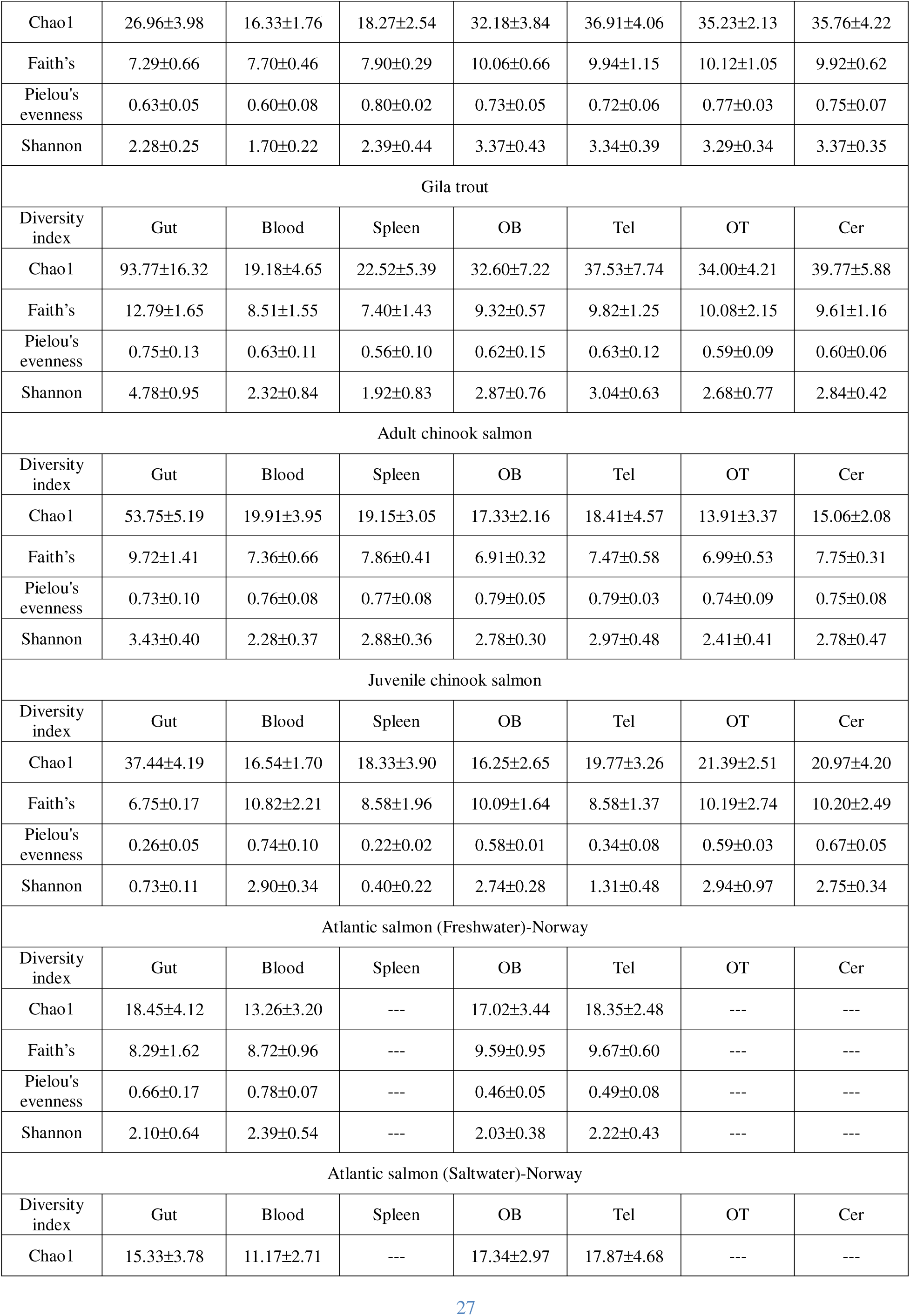
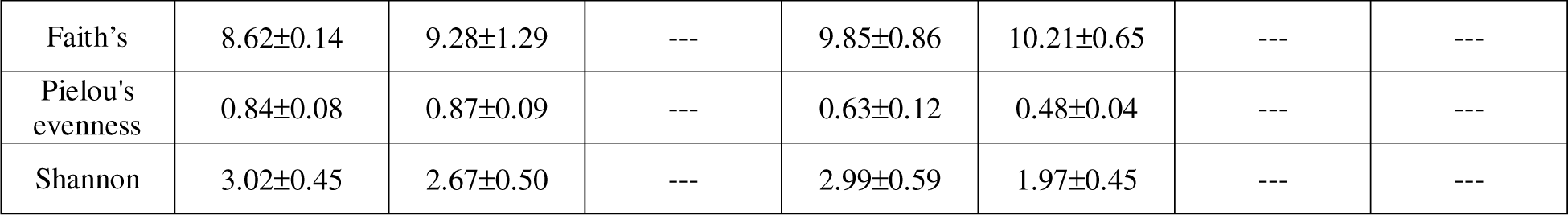
Diversity indexes ± SD for all geographically covered salmonid species.

**Table S3.** Laboratory rainbow trout brain and spleen microbial community diversity proportions predicted to originate from various sources using SourceTracker2 (Mean ± SD).

| Source<br>Tissue |  | Blood | Gut | Water | Unknown |
| --- | --- | --- | --- | --- | --- |
| Diversity proportion | Spleen | 11.11±2.80 | 27.77±6.33 | 3.45±0.60 | 57.65±6.76 |
|  | OB | 12.52±4.00 | 12.50±8.42 | 2.35± 0.96 | 72.64± 5.77** |
|  | Tel | 16.07±4.11 | 15.92±9.07 | 2.88±1.16 | 65.11±7.14 |
|  | OT | 18.18±3.70 | 22.72±8.72 | 1.91±1.07 | 57.17±5.74 |
|  | Cer | 21.21±3.21 | 6.06±6.44 | 4.77±1.15* | 67.95±6.37 |

\* P-value= 0.0137 \*\* P-value=0.0066

**Table S4.** Summary of *Agrobacterium* sp. and *Plesiomonas* sp. isolates selected for whole genome sequencing. (Sh=short genome; L=long genome).

| Bacterial Isolate | Fish number1 |  |  | Fish Number2 |  |  |
| --- | --- | --- | --- | --- | --- | --- |
| Isolation source | Gut | Blood | Brain | Gut | Blood | Brain |
| *Agrobacterium* sp. | 3 (Type strain) | 3 (Type strain) | 2 (exoP mutant) | 2 (Type strain) | 2 (Type strain) | 2 (exoP mutant) |
| *Plesiomonas* sp. | 3 (1Sh+2L) | 2(1Sh+1L) | 3 (3L) | 3 (1Sh+1L) | 2(1Sh+1L) | 3 (3L) |

**Table S5.**
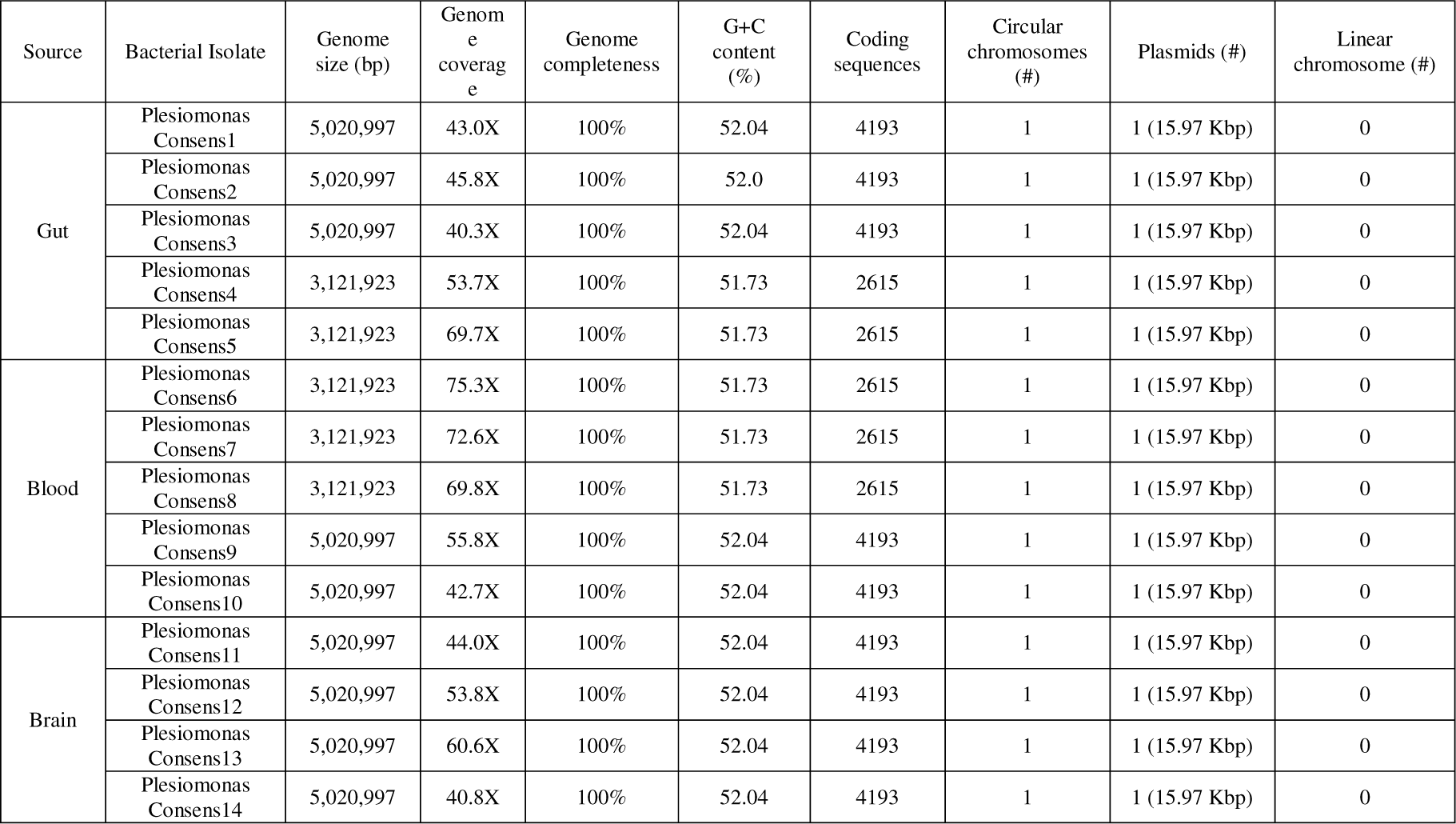
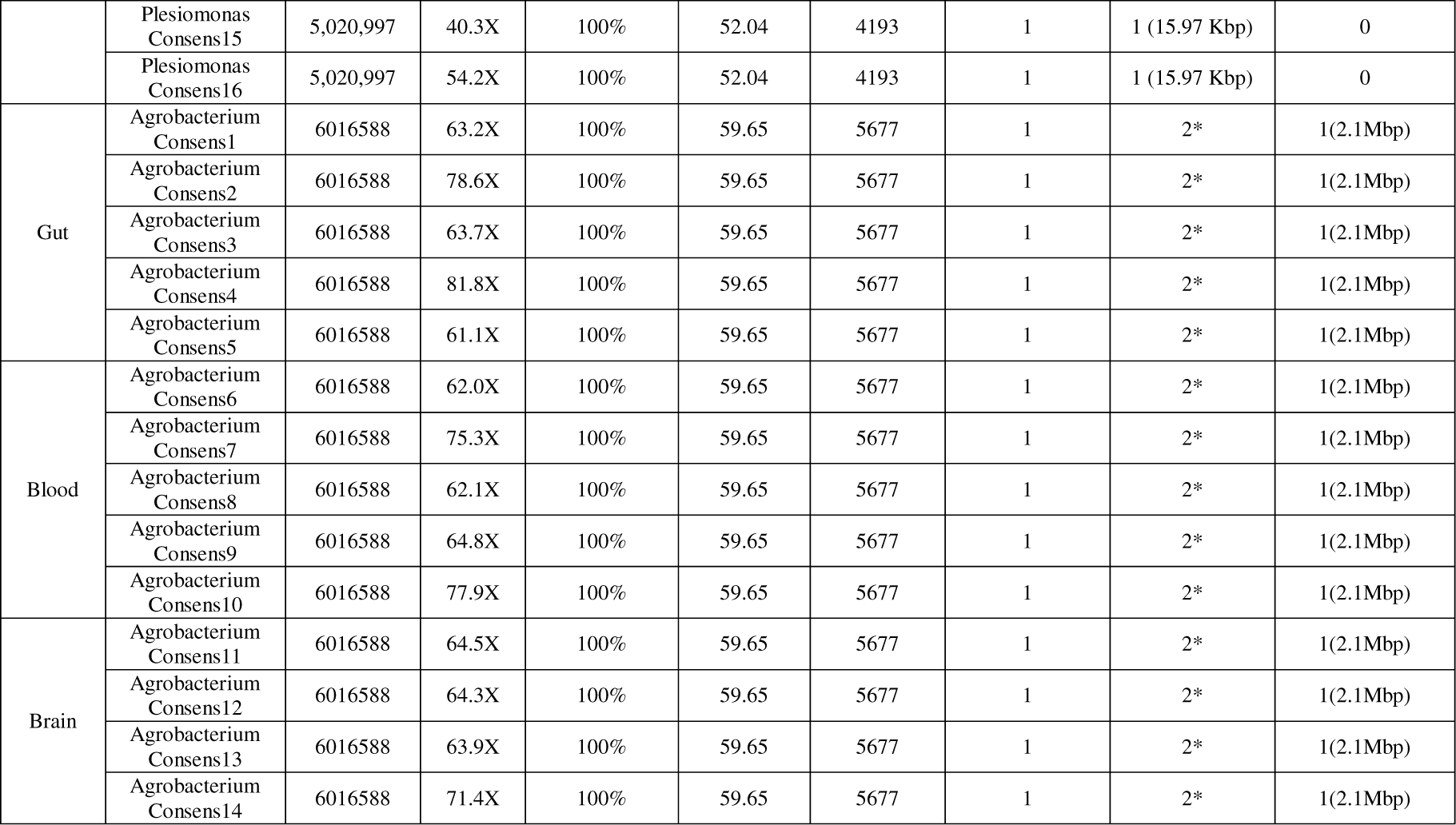
Summary of *Plesiomonas* sp. and *Agrobacterium* sp. genomes sequenced from rainbow trout gut, blood, and brain in this study.

| Source | Bacterial Isolate | Genome size (bp) | Genome coverage | Genome completeness | G+C content (%) | Coding sequences | Circular chromosomes (#) | Plasmids (#) | Linear chromosome (#) |
| --- | --- | --- | --- | --- | --- | --- | --- | --- | --- |
| Gut | *Plesiomonas* Consens1 | 5,020,997 | 43.0X | 100% | 52.04 | 4193 | 1 | 1 (15.97 Kbp) | 0 |
|  | *Plesiomonas* Consens2 | 5,020,997 | 45.8X | 100% | 52.0 | 4193 | 1 | 1 (15.97 Kbp) | 0 |
|  | *Plesiomonas* Consens3 | 5,020,997 | 40.3X | 100% | 52.04 | 4193 | 1 | 1 (15.97 Kbp) | 0 |
|  | *Plesiomonas* Consens4 | 3,121,923 | 53.7X | 100% | 51.73 | 2615 | 1 | 1 (15.97 Kbp) | 0 |
|  | *Plesiomonas* Consens5 | 3,121,923 | 69.7X | 100% | 51.73 | 2615 | 1 | 1 (15.97 Kbp) | 0 |
| Blood | *Plesiomonas* Consens6 | 3,121,923 | 75.3X | 100% | 51.73 | 2615 | 1 | 1 (15.97 Kbp) | 0 |
|  | *Plesiomonas* Consens7 | 3,121,923 | 72.6X | 100% | 51.73 | 2615 | 1 | 1 (15.97 Kbp) | 0 |
|  | *Plesiomonas* Consens8 | 3,121,923 | 69.8X | 100% | 51.73 | 2615 | 1 | 1 (15.97 Kbp) | 0 |
|  | *Plesiomonas* Consens9 | 5,020,997 | 55.8X | 100% | 52.04 | 4193 | 1 | 1 (15.97 Kbp) | 0 |
|  | *Plesiomonas* Consens10 | 5,020,997 | 42.7X | 100% | 52.04 | 4193 | 1 | 1 (15.97 Kbp) | 0 |
| Brain | *Plesiomonas* Consens11 | 5,020,997 | 44.0X | 100% | 52.04 | 4193 | 1 | 1 (15.97 Kbp) | 0 |
|  | *Plesiomonas* Consens12 | 5,020,997 | 53.8X | 100% | 52.04 | 4193 | 1 | 1 (15.97 Kbp) | 0 |
|  | *Plesiomonas* Consens13 | 5,020,997 | 60.6X | 100% | 52.04 | 4193 | 1 | 1 (15.97 Kbp) | 0 |
|  | *Plesiomonas* Consens14 | 5,020,997 | 40.8X | 100% | 52.04 | 4193 | 1 | 1 (15.97 Kbp) | 0 |
|  | Plesiomonas Consens15 | 5,020,997 | 40.3X | 100% | 52.04 | 4193 | 1 | 1 (15.97 Kbp) | 0 |
|  | Plesiomonas Consens16 | 5,020,997 | 54.2X | 100% | 52.04 | 4193 | 1 | 1 (15.97 Kbp) | 0 |
| Gut | Agrobacterium Consens1 | 6016588 | 63.2X | 100% | 59.65 | 5677 | 1 | 2* | 1(2.1Mbp) |
|  | Agrobacterium Consens2 | 6016588 | 78.6X | 100% | 59.65 | 5677 | 1 | 2* | 1(2.1Mbp) |
|  | Agrobacterium Consens3 | 6016588 | 63.7X | 100% | 59.65 | 5677 | 1 | 2* | 1(2.1Mbp) |
|  | Agrobacterium Consens4 | 6016588 | 81.8X | 100% | 59.65 | 5677 | 1 | 2* | 1(2.1Mbp) |
|  | Agrobacterium Consens5 | 6016588 | 61.1X | 100% | 59.65 | 5677 | 1 | 2* | 1(2.1Mbp) |
| Blood | Agrobacterium Consens6 | 6016588 | 62.0X | 100% | 59.65 | 5677 | 1 | 2* | 1(2.1Mbp) |
|  | Agrobacterium Consens7 | 6016588 | 75.3X | 100% | 59.65 | 5677 | 1 | 2* | 1(2.1Mbp) |
|  | Agrobacterium Consens8 | 6016588 | 62.1X | 100% | 59.65 | 5677 | 1 | 2* | 1(2.1Mbp) |
|  | Agrobacterium Consens9 | 6016588 | 64.8X | 100% | 59.65 | 5677 | 1 | 2* | 1(2.1Mbp) |
|  | Agrobacterium Consens10 | 6016588 | 77.9X | 100% | 59.65 | 5677 | 1 | 2* | 1(2.1Mbp) |
| Brain | Agrobacterium Consens11 | 6016588 | 64.5X | 100% | 59.65 | 5677 | 1 | 2* | 1(2.1Mbp) |
|  | Agrobacterium Consens12 | 6016588 | 64.3X | 100% | 59.65 | 5677 | 1 | 2* | 1(2.1Mbp) |
|  | Agrobacterium Consens13 | 6016588 | 63.9X | 100% | 59.65 | 5677 | 1 | 2* | 1(2.1Mbp) |
|  | Agrobacterium Consens14 | 6016588 | 71.4X | 100% | 59.65 | 5677 | 1 | 2* | 1(2.1Mbp) |

**Table S6.** Summary of module completeness for 39 significantly different KEGG pathways between short genome and long genome *Plesiomonas* sp. performed in Anvi’o.

| Module name | MC (Short genome) | MC (Long genome) | Adjusted P-value |
| --- | --- | --- | --- |
| PpkA biosynthesis | 0.45 | 0.80 | 1.29E-04 |
| Reductive citrate cycle | 0.17 | 0.83 | 1.32E-04 |
| C4-dicarboxylic acid cycle | 0.06 | 0.53 | 2.67E-04 |
| NAD cycle | 0.63 | 0.88 | 3.10E-04 |
| Acylglycerol degradation | 0.80 | 1.00 | 3.92E-04 |
| Methionine degradation | 0.20 | 0.80 | 4.59E-04 |
| YscQL biosynthesis | 0.60 | 1.00 | 7.80E-04 |
| Glycogen degradation | 0.33 | 0.67 | 1.10E-03 |
| YscN biosynthesis | 0.60 | 0.90 | 1.18E-03 |
| Putrescine => Spermidine | 0.00 | 0.75 | 1.20E-03 |
| D-galactonate degradation | 0.11 | 0.28 | 1.24E-03 |
| YopD biosynthesis | 0.65 | 1.00 | 1.29E-03 |
| Denitrification | 0.50 | 1.00 | 1.30E-03 |
| YscW biosynthesis | 0.45 | 0.90 | 1.46E-03 |
| Ethylmalonyl pathway | 0.20 | 0.40 | 1.48E-03 |
| D-Galacturonate degradation | 0.75 | 1.00 | 1.52E-03 |
| TCA cycle | 0.88 | 1.00 | 1.53E-03 |
| Methylaspartate cycle | 0.67 | 1.00 | 1.58E-03 |
| Lip biosynthesis | 0.25 | 0.90 | 1.59E-03 |
| Arginine => Ornithine | 0.00 | 1.00 | 1.68E-03 |
| DotU biosynthesis | 0.45 | 0.80 | 1.70E-03 |
| ICMF biosynthesis | 0.50 | 0.73 | 1.71E-03 |
| ClpV biosynthesis | 0.40 | 0.75 | 1.72E-03 |
| Adenine ribonucleotide biosynthesis | 0.60 | 0.80 | 1.78E-03 |
| Galactose degradation | 0.22 | 0.83 | 1.79E-03 |
| Reductive citrate cycle | 0.10 | 0.35 | 1.97E-03 |
| YscC biosynthesis | 0.33 | 1.00 | 2.10E-03 |
| YscJ biosynthesis | 0.45 | 1.00 | 2.13E-03 |
| Dicarboxylate cycle | 0.43 | 1.00 | 2.14E-03 |
| Ketone body biosynthesis | 0.52 | 0.69 | 2.15E-03 |
| Glyoxylate cycle | 0.72 | 0.83 | 2.18E-03 |
| nitrate reduction | 0.50 | 0.75 | 2.20E-03 |
| Hydroxypropionate cycle | 0.09 | 0.82 | 2.29E-03 |
| Incomplete reductive citrate | 0.33 | 0.56 | 2.39E-03 |
| Ascorbate biosynthesis | 0.60 | 0.80 | 2.44E-03 |
| CAM (Crassulacean acid metabolism) | 0.44 | 0.56 | 2.53E-03 |
| Leucine degradation | 0.20 | 0.80 | 2.58E-03 |
| Glycogen biosynthesis | 0.50 | 0.67 | 2.63E-03 |

