## Supplemental File 2 for "A niche-adapted brain microbiome in salmonids at homeostasis"

**SPEC**

>Consens1

MKTLKIAASDSVLSCFDTEREITNVHTTDFSDIAAIVVSVQDIHDGILAKIHATGLSIPTFAAVCCEEELSAEVLPQLTGVFELCGENTDFYGKQLESAAQRYEENLLPPFFNTLKQYVEMGNSTFACPGHQGGQFFRKHPAGRQFFEFFGETLFRADMCNADVKLGDLLIHEGAPCDAQKHAAKVYNADKTYFVLNGTSASNKVATNALLARGDLVLFDRNNHKSNHHGALLQAGATPVYLETARNPFGFIGGIDAHCFSEKYLRDEIRKVAPEKAEAKRPFRLAIIQLGTYDGTIYNARQVVDKIGHLCDYILFDSAWVGYEQFIPMMKDCSPLLLDLNENDPGIIVTQSVHKQQAGFSQTSQIHKKDSHIKGQDRYCNHKRFNNAFMMHASTSPFYPLFAALDVNAKMHEGESGRRLWMECVKNGIEARKLLLETCSMIKPFVPVTVDGRKWQDHDTEVMANDLRFFNFVPSEKWHGFEGYQEHQYFVDPCKFMLTTPGIDAATGEYTEFGIPATILANFLRENGIIPEKCDLNSILFLLTPAEDMAKLQHLIAQIARFERYVAEDAPLSEVLPNVYRSNEARYRGYTIRQLCQEMHNLYVSYDVKQLQKEMFRQAHFPKAVMNPQDANIEFIRGNVELVPLCKAEGRIAAEGALPYPPGVLCVVPGEVWGGAVQRYFLALEEGINLLPGFSPELQGVYIQEEDDGTKRAYGYVMKQ

>Consens2

MKTLKIAASDSVLSCFDTEREITNVHTTDFSDIAAIVVSVQDIHDGILAKIHATGLSIPTFAAVCCEEELSAEVLPQLTGVFELCGENTDFYGKQLESAAQRYEENLLPPFFNTLKQYVEMGNSTFACPGHQGGQFFRKHPAGRQFFEFFGETLFRADMCNADVKLGDLLIHEGAPCDAQKHAAKVYNADKTYFVLNGTSASNKVATNALLARGDLVLFDRNNHKSNHHGALLQAGATPVYLETARNPFGFIGGIDAHCFSEKYLRDEIRKVAPEKAEAKRPFRLAIIQLGTYDGTIYNARQVVDKIGHLCDYILFDSAWVGYEQFIPMMKDCSPLLLDLNENDPGIIVTQSVHKQQAGFSQTSQIHKKDSHIKGQDRYCNHKRFNNAFMMHASTSPFYPLFAALDVNAKMHEGESGRRLWMECVKNGIEARKLLLETCSMIKPFVPVTVDGRKWQDHDTEVMANDLRFFNFVPSEKWHGFEGYQEHQYFVDPCKFMLTTPGIDAATGEYTEFGIPATILANFLRENGIIPEKCDLNSILFLLTPAEDMAKLQHLIAQIARFERYVAEDAPLSEVLPNVYRSNEARYRGYTIRQLCQEMHNLYVSYDVKQLQKEMFRQAHFPKAVMNPQDANIEFIRGNVELVPLCKAEGRIAAEGALPYPPGVLCVVPGEVWGGAVQRYFLALEEGINLLPGFSPELQGVYIQEEDDGTKRAYGYVMKQ

>Consens3

MKTLKIAASDSVLSCFDTEREITNVHTTDFSDIAAIVVSVQDIHDGILAKIHATGLSIPTFAAVCCEEELSAEVLPQLTGVFELCGENTDFYGKQLESAAQRYEENLLPPFFNTLKQYVEMGNSTFACPGHQGGQFFRKHPAGRQFFEFFGETLFRADMCNADVKLGDLLIHEGAPCDAQKHAAKVYNADKTYFVLNGTSASNKVATNALLARGDLVLFDRNNHKSNHHGALLQAGATPVYLETARNPFGFIGGIDAHCFSEKYLRDEIRKVAPEKAEAKRPFRLAIIQLGTYDGTIYNARQVVDKIGHLCDYILFDSAWVGYEQFIPMMKDCSPLLLDLNENDPGIIVTQSVHKQQAGFSQTSQIHKKDSHIKGQDRYCNHKRFNNAFMMHASTSPFYPLFAALDVNAKMHEGESGRRLWMECVKNGIEARKLLLETCSMIKPFVPVTVDGRKWQDHDTEVMANDLRFFNFVPSEKWHGFEGYQEHQYFVDPCKFMLTTPGIDAATGEYTEFGIPATILANFLRENGIIPEKCDLNSILFLLTPAEDMAKLQHLIAQIARFERYVAEDAPLSEVLPNVYRSNEARYRGYTIRQLCQEMHNLYVSYDVKQLQKEMFRQAHFPKAVMNPQDANIEFIRGNVELVPLCKAEGRIAAEGALPYPPGVLCVVPGEVWGGAVQRYFLALEEGINLLPGFSPELQGVYIQEEDDGTKRAYGYVMKQ

>Consens4

MKTLKIAASDSVLSCFDTEREITNVHTTDFSDIAAIVVSVQDIHDGILAKIHATGLSIPTFAAVCCEEELSAEVLPQLTGVFELCGENTDFYGKQLESAAQRYEENLLPPFFNTLKQYVEMGNSTFACPGHQGGQFFRKHPAGRQFFEFFGETLFRADMCNADVKLGDLLIHEGAPCDAQKHAAKVYNADKTYFVLNGTSASNKVATNALLARGDLVLFDRNNHKSNHHGALLQAGATPVYLETARNPFGFIGGIDAHCFSEKYLRDEIRKVAPEKAEAKRPFRLAIIQLGTYDGTIYNARQVVDKIGHLCDYILFDSAWVGYEQFIPMMKDCSPLLLDLNENDPGIIVTQSVHKQQAGFSQTSQIHKKDSHIKGQDRYCNHKRFNNAFMMHASTSPFYPLFAALDVNAKMHEGESGRRLWMECVKNGIEARKLLLETCSMIKPFVPVTVDGRKWQDHDTEVMANDLRFFNFVPSEKWHGFEGYQEHQYFVDPCKFMLTTPGIDAATGEYTEFGIPATILANFLRENGIIPEKCDLNSILFLLTPAEDMAKLQHLIAQIARFERYVAEDAPLSEVLPNVYRSNEARYRGYTIRQLCQEMHNLYVSYDVKQLQKEMFRQAHFPKAVMNPQDANIEFIRGNVELVPLCKAEGRIAAEGALPYPPGVLCVVPGEVWGGAVQRYFLALEEGINLLPGFSPELQGVYIQEEDDGTKRAYGYVMKQ

>Consens5

MKTLKIAASDSVLSCFDTEREITNVHTTDFSDIAAIVVSVQDIHDGILAKIHATGLSIPTFAAVCCEEELSAEVLPQLTGVFELCGENTDFYGKQLESAAQRYEENLLPPFFNTLKQYVEMGNSTFACPGHQGGQFFRKHPAGRQFFEFFGETLFRADMCNADVKLGDLLIHEGAPCDAQKHAAKVYNADKTYFVLNGTSASNKVATNALLARGDLVLFDRNNHKSNHHGALLQAGATPVYLETARNPFGFIGGIDAHCFSEKYLRDEIRKVAPEKAEAKRPFRLAIIQLGTYDGTIYNARQVVDKIGHLCDYILFDSAWVGYEQFIPMMKDCSPLLLDLNENDPGIIVTQSVHKQQAGFSQTSQIHKKDSHIKGQDRYCNHKRFNNAFMMHASTSPFYPLFAALDVNAKMHEGESGRRLWMECVKNGIEARKLLLETCSMIKPFVPVTVDGRKWQDHDTEVMANDLRFFNFVPSEKWHGFEGYQEHQYFVDPCKFMLTTPGIDAATGEYTEFGIPATILANFLRENGIIPEKCDLNSILFLLTPAEDMAKLQHLIAQIARFERYVAEDAPLSEVLPNVYRSNEARYRGYTIRQLCQEMHNLYVSYDVKQLQKEMFRQAHFPKAVMNPQDANIEFIRGNVELVPLCKAEGRIAAEGALPYPPGVLCVVPGEVWGGAVQRYFLALEEGINLLPGFSPELQGVYIQEEDDGTKRAYGYVMKQ

>Consens6

MKTLKIAASDSVLSCFDTEREITNVHTTDFSDIAAIVVSVQDIHDGILAKIHATGLSIPTFAAVCCEEELSAEVLPQLTGVFELCGENTDFYGKQLESAAQRYEENLLPPFFNTLKQYVEMGNSTFACPGHQGGQFFRKHPAGRQFFEFFGETLFRADMCNADVKLGDLLIHEGAPCDAQKHAAKVYNADKTYFVLNGTSASNKVATNALLARGDLVLFDRNNHKSNHHGALLQAGATPVYLETARNPFGFIGGIDAHCFSEKYLRDEIRKVAPEKAEAKRPFRLAIIQLGTYDGTIYNARQVVDKIGHLCDYILFDSAWVGYEQFIPMMKDCSPLLLDLNENDPGIIVTQSVHKQQAGFSQTSQIHKKDSHIKGQDRYCNHKRFNNAFMMHASTSPFYPLFAALDVNAKMHEGESGRRLWMECVKNGIEARKLLLETCSMIKPFVPVTVDGRKWQDHDTEVMANDLRFFNFVPSEKWHGFEGYQEHQYFVDPCKFMLTTPGIDAATGEYTEFGIPATILANFLRENGIIPEKCDLNSILFLLTPAEDMAKLQHLIAQIARFERYVAEDAPLSEVLPNVYRSNEARYRGYTIRQLCQEMHNLYVSYDVKQLQKEMFRQAHFPKAVMNPQDANIEFIRGNVELVPLCKAEGRIAAEGALPYPPGVLCVVPGEVWGGAVQRYFLALEEGINLLPGFSPELQGVYIQEEDDGTKRAYGYVMKQ

>Consens7

MKTLKIAASDSVLSCFDTEREITNVHTTDFSDIAAIVVSVQDIHDGILAKIHATGLSIPTFAAVCCEEELSAEVLPQLTGVFELCGENTDFYGKQLESAAQRYEENLLPPFFNTLKQYVEMGNSTFACPGHQGGQFFRKHPAGRQFFEFFGETLFRADMCNADVKLGDLLIHEGAPCDAQKHAAKVYNADKTYFVLNGTSASNKVATNALLARGDLVLFDRNNHKSNHHGALLQAGATPVYLETARNPFGFIGGIDAHCFSEKYLRDEIRKVAPEKAEAKRPFRLAIIQLGTYDGTIYNARQVVDKIGHLCDYILFDSAWVGYEQFIPMMKDCSPLLLDLNENDPGIIVTQSVHKQQAGFSQTSQIHKKDSHIKGQDRYCNHKRFNNAFMMHASTSPFYPLFAALDVNAKMHEGESGRRLWMECVKNGIEARKLLLETCSMIKPFVPVTVDGRKWQDHDTEVMANDLRFFNFVPSEKWHGFEGYQEHQYFVDPCKFMLTTPGIDAATGEYTEFGIPATILANFLRENGIIPEKCDLNSILFLLTPAEDMAKLQHLIAQIARFERYVAEDAPLSEVLPNVYRSNEARYRGYTIRQLCQEMHNLYVSYDVKQLQKEMFRQAHFPKAVMNPQDANIEFIRGNVELVPLCKAEGRIAAEGALPYPPGVLCVVPGEVWGGAVQRYFLALEEGINLLPGFSPELQGVYIQEEDDGTKRAYGYVMKQ

>Consens8

MKTLKIAASDSVLSCFDTEREITNVHTTDFSDIAAIVVSVQDIHDGILAKIHATGLSIPTFAAVCCEEELSAEVLPQLTGVFELCGENTDFYGKQLESAAQRYEENLLPPFFNTLKQYVEMGNSTFACPGHQGGQFFRKHPAGRQFFEFFGETLFRADMCNADVKLGDLLIHEGAPCDAQKHAAKVYNADKTYFVLNGTSASNKVATNALLARGDLVLFDRNNHKSNHHGALLQAGATPVYLETARNPFGFIGGIDAHCFSEKYLRDEIRKVAPEKAEAKRPFRLAIIQLGTYDGTIYNARQVVDKIGHLCDYILFDSAWVGYEQFIPMMKDCSPLLLDLNENDPGIIVTQSVHKQQAGFSQTSQIHKKDSHIKGQDRYCNHKRFNNAFMMHASTSPFYPLFAALDVNAKMHEGESGRRLWMECVKNGIEARKLLLETCSMIKPFVPVTVDGRKWQDHDTEVMANDLRFFNFVPSEKWHGFEGYQEHQYFVDPCKFMLTTPGIDAATGEYTEFGIPATILANFLRENGIIPEKCDLNSILFLLTPAEDMAKLQHLIAQIARFERYVAEDAPLSEVLPNVYRSNEARYRGYTIRQLCQEMHNLYVSYDVKQLQKEMFRQAHFPKAVMNPQDANIEFIRGNVELVPLCKAEGRIAAEGALPYPPGVLCVVPGEVWGGAVQRYFLALEEGINLLPGFSPELQGVYIQEEDDGTKRAYGYVMKQ

>Consens9

MKTLKIAASDSVLSCFDTEREITNVHTTDFSDIAAIVVSVQDIHDGILAKIHATGLSIPTFAAVCCEEELSAEVLPQLTGVFELCGENTDFYGKQLESAAQRYEENLLPPFFNTLKQYVEMGNSTFACPGHQGGQFFRKHPAGRQFFEFFGETLFRADMCNADVKLGDLLIHEGAPCDAQKHAAKVYNADKTYFVLNGTSASNKVATNALLARGDLVLFDRNNHKSNHHGALLQAGATPVYLETARNPFGFIGGIDAHCFSEKYLRDEIRKVAPEKAEAKRPFRLAIIQLGTYDGTIYNARQVVDKIGHLCDYILFDSAWVGYEQFIPMMKDCSPLLLDLNENDPGIIVTQSVHKQQAGFSQTSQIHKKDSHIKGQDRYCNHKRFNNAFMMHASTSPFYPLFAALDVNAKMHEGESGRRLWMECVKNGIEARKLLLETCSMIKPFVPVTVDGRKWQDHDTEVMANDLRFFNFVPSEKWHGFEGYQEHQYFVDPCKFMLTTPGIDAATGEYTEFGIPATILANFLRENGIIPEKCDLNSILFLLTPAEDMAKLQHLIAQIARFERYVAEDAPLSEVLPNVYRSNEARYRGYTIRQLCQEMHNLYVSYDVKQLQKEMFRQAHFPKAVMNPQDANIEFIRGNVELVPLCKAEGRIAAEGALPYPPGVLCVVPGEVWGGAVQRYFLALEEGINLLPGFSPELQGVYIQEEDDGTKRAYGYVMKQ

>Consens10

MKTLKIAASDSVLSCFDTEREITNVHTTDFSDIAAIVVSVQDIHDGILAKIHATGLSIPTFAAVCCEEELSAEVLPQLTGVFELCGENTDFYGKQLESAAQRYEENLLPPFFNTLKQYVEMGNSTFACPGHQGGQFFRKHPAGRQFFEFFGETLFRADMCNADVKLGDLLIHEGAPCDAQKHAAKVYNADKTYFVLNGTSASNKVATNALLARGDLVLFDRNNHKSNHHGALLQAGATPVYLETARNPFGFIGGIDAHCFSEKYLRDEIRKVAPEKAEAKRPFRLAIIQLGTYDGTIYNARQVVDKIGHLCDYILFDSAWVGYEQFIPMMKDCSPLLLDLNENDPGIIVTQSVHKQQAGFSQTSQIHKKDSHIKGQDRYCNHKRFNNAFMMHASTSPFYPLFAALDVNAKMHEGESGRRLWMECVKNGIEARKLLLETCSMIKPFVPVTVDGRKWQDHDTEVMANDLRFFNFVPSEKWHGFEGYQEHQYFVDPCKFMLTTPGIDAATGEYTEFGIPATILANFLRENGIIPEKCDLNSILFLLTPAEDMAKLQHLIAQIARFERYVAEDAPLSEVLPNVYRSNEARYRGYTIRQLCQEMHNLYVSYDVKQLQKEMFRQAHFPKAVMNPQDANIEFIRGNVELVPLCKAEGRIAAEGALPYPPGVLCVVPGEVWGGAVQRYFLALEEGINLLPGFSPELQGVYIQEEDDGTKRAYGYVMKQ

>Consens11

MKTLKIAASDSVLSCFDTEREITNVHTTDFSDIAAIVVSVQDIHDGILAKIHATGLSIPTFAAVCCEEELSAEVLPQLTGVFELCGENTDFYGKQLESAAQRYEENLLPPFFNTLKQYVEMGNSTFACPGHQGGQFFRKHPAGRQFFEFFGETLFRADMCNADVKLGDLLIHEGAPCDAQKHAAKVYNADKTYFVLNGTSASNKVATNALLARGDLVLFDRNNHKSNHHGALLQAGATPVYLETARNPFGFIGGIDAHCFSEKYLRDEIRKVAPEKAEAKRPFRLAIIQLGTYDGTIYNARQVVDKIGHLCDYILFDSAWVGYEQFIPMMKDCSPLLLDLNENDPGIIVTQSVHKQQAGFSQTSQIHKKDSHIKGQDRYCNHKRFNNAFMMHASTSPFYPLFAALDVNAKMHEGESGRRLWMECVKNGIEARKLLLETCSMIKPFVPVTVDGRKWQDHDTEVMANDLRFFNFVPSEKWHGFEGYQEHQYFVDPCKFMLTTPGIDAATGEYTEFGIPATILANFLRENGIIPEKCDLNSILFLLTPAEDMAKLQHLIAQIARFERYVAEDAPLSEVLPNVYRSNEARYRGYTIRQLCQEMHNLYVSYDVKQLQKEMFRQAHFPKAVMNPQDANIEFIRGNVELVPLCKAEGRIAAEGALPYPPGVLCVVPGEVWGGAVQRYFLALEEGINLLPGFSPELQGVYIQEEDDGTKRAYGYVMKQ

>Consens12

MKTLKIAASDSVLSCFDTEREITNVHTTDFSDIAAIVVSVQDIHDGILAKIHATGLSIPTFAAVCCEEELSAEVLPQLTGVFELCGENTDFYGKQLESAAQRYEENLLPPFFNTLKQYVEMGNSTFACPGHQGGQFFRKHPAGRQFFEFFGETLFRADMCNADVKLGDLLIHEGAPCDAQKHAAKVYNADKTYFVLNGTSASNKVATNALLARGDLVLFDRNNHKSNHHGALLQAGATPVYLETARNPFGFIGGIDAHCFSEKYLRDEIRKVAPEKAEAKRPFRLAIIQLGTYDGTIYNARQVVDKIGHLCDYILFDSAWVGYEQFIPMMKDCSPLLLDLNENDPGIIVTQSVHKQQAGFSQTSQIHKKDSHIKGQDRYCNHKRFNNAFMMHASTSPFYPLFAALDVNAKMHEGESGRRLWMECVKNGIEARKLLLETCSMIKPFVPVTVDGRKWQDHDTEVMANDLRFFNFVPSEKWHGFEGYQEHQYFVDPCKFMLTTPGIDAATGEYTEFGIPATILANFLRENGIIPEKCDLNSILFLLTPAEDMAKLQHLIAQIARFERYVAEDAPLSEVLPNVYRSNEARYRGYTIRQLCQEMHNLYVSYDVKQLQKEMFRQAHFPKAVMNPQDANIEFIRGNVELVPLCKAEGRIAAEGALPYPPGVLCVVPGEVWGGAVQRYFLALEEGINLLPGFSPELQGVYIQEEDDGTKRAYGYVMKQ

>Consens13

MKTLKIAASDSVLSCFDTEREITNVHTTDFSDIAAIVVSVQDIHDGILAKIHATGLSIPTFAAVCCEEELSAEVLPQLTGVFELCGENTDFYGKQLESAAQRYEENLLPPFFNTLKQYVEMGNSTFACPGHQGGQFFRKHPAGRQFFEFFGETLFRADMCNADVKLGDLLIHEGAPCDAQKHAAKVYNADKTYFVLNGTSASNKVATNALLARGDLVLFDRNNHKSNHHGALLQAGATPVYLETARNPFGFIGGIDAHCFSEKYLRDEIRKVAPEKAEAKRPFRLAIIQLGTYDGTIYNARQVVDKIGHLCDYILFDSAWVGYEQFIPMMKDCSPLLLDLNENDPGIIVTQSVHKQQAGFSQTSQIHKKDSHIKGQDRYCNHKRFNNAFMMHASTSPFYPLFAALDVNAKMHEGESGRRLWMECVKNGIEARKLLLETCSMIKPFVPVTVDGRKWQDHDTEVMANDLRFFNFVPSEKWHGFEGYQEHQYFVDPCKFMLTTPGIDAATGEYTEFGIPATILANFLRENGIIPEKCDLNSILFLLTPAEDMAKLQHLIAQIARFERYVAEDAPLSEVLPNVYRSNEARYRGYTIRQLCQEMHNLYVSYDVKQLQKEMFRQAHFPKAVMNPQDANIEFIRGNVELVPLCKAEGRIAAEGALPYPPGVLCVVPGEVWGGAVQRYFLALEEGINLLPGFSPELQGVYIQEEDDGTKRAYGYVMKQ

>Consens14

MKTLKIAASDSVLSCFDTEREITNVHTTDFSDIAAIVVSVQDIHDGILAKIHATGLSIPTFAAVCCEEELSAEVLPQLTGVFELCGENTDFYGKQLESAAQRYEENLLPPFFNTLKQYVEMGNSTFACPGHQGGQFFRKHPAGRQFFEFFGETLFRADMCNADVKLGDLLIHEGAPCDAQKHAAKVYNADKTYFVLNGTSASNKVATNALLARGDLVLFDRNNHKSNHHGALLQAGATPVYLETARNPFGFIGGIDAHCFSEKYLRDEIRKVAPEKAEAKRPFRLAIIQLGTYDGTIYNARQVVDKIGHLCDYILFDSAWVGYEQFIPMMKDCSPLLLDLNENDPGIIVTQSVHKQQAGFSQTSQIHKKDSHIKGQDRYCNHKRFNNAFMMHASTSPFYPLFAALDVNAKMHEGESGRRLWMECVKNGIEARKLLLETCSMIKPFVPVTVDGRKWQDHDTEVMANDLRFFNFVPSEKWHGFEGYQEHQYFVDPCKFMLTTPGIDAATGEYTEFGIPATILANFLRENGIIPEKCDLNSILFLLTPAEDMAKLQHLIAQIARFERYVAEDAPLSEVLPNVYRSNEARYRGYTIRQLCQEMHNLYVSYDVKQLQKEMFRQAHFPKAVMNPQDANIEFIRGNVELVPLCKAEGRIAAEGALPYPPGVLCVVPGEVWGGAVQRYFLALEEGINLLPGFSPELQGVYIQEEDDGTKRAYGYVMKQ

>Consens15

MKTLKIAASDSVLSCFDTEREITNVHTTDFSDIAAIVVSVQDIHDGILAKIHATGLSIPTFAAVCCEEELSAEVLPQLTGVFELCGENTDFYGKQLESAAQRYEENLLPPFFNTLKQYVEMGNSTFACPGHQGGQFFRKHPAGRQFFEFFGETLFRADMCNADVKLGDLLIHEGAPCDAQKHAAKVYNADKTYFVLNGTSASNKVATNALLARGDLVLFDRNNHKSNHHGALLQAGATPVYLETARNPFGFIGGIDAHCFSEKYLRDEIRKVAPEKAEAKRPFRLAIIQLGTYDGTIYNARQVVDKIGHLCDYILFDSAWVGYEQFIPMMKDCSPLLLDLNENDPGIIVTQSVHKQQAGFSQTSQIHKKDSHIKGQDRYCNHKRFNNAFMMHASTSPFYPLFAALDVNAKMHEGESGRRLWMECVKNGIEARKLLLETCSMIKPFVPVTVDGRKWQDHDTEVMANDLRFFNFVPSEKWHGFEGYQEHQYFVDPCKFMLTTPGIDAATGEYTEFGIPATILANFLRENGIIPEKCDLNSILFLLTPAEDMAKLQHLIAQIARFERYVAEDAPLSEVLPNVYRSNEARYRGYTIRQLCQEMHNLYVSYDVKQLQKEMFRQAHFPKAVMNPQDANIEFIRGNVELVPLCKAEGRIAAEGALPYPPGVLCVVPGEVWGGAVQRYFLALEEGINLLPGFSPELQGVYIQEEDDGTKRAYGYVMKQ

>Consens16

MKTLKIAASDSVLSCFDTEREITNVHTTDFSDIAAIVVSVQDIHDGILAKIHATGLSIPTFAAVCCEEELSAEVLPQLTGVFELCGENTDFYGKQLESAAQRYEENLLPPFFNTLKQYVEMGNSTFACPGHQGGQFFRKHPAGRQFFEFFGETLFRADMCNADVKLGDLLIHEGAPCDAQKHAAKVYNADKTYFVLNGTSASNKVATNALLARGDLVLFDRNNHKSNHHGALLQAGATPVYLETARNPFGFIGGIDAHCFSEKYLRDEIRKVAPEKAEAKRPFRLAIIQLGTYDGTIYNARQVVDKIGHLCDYILFDSAWVGYEQFIPMMKDCSPLLLDLNENDPGIIVTQSVHKQQAGFSQTSQIHKKDSHIKGQDRYCNHKRFNNAFMMHASTSPFYPLFAALDVNAKMHEGESGRRLWMECVKNGIEARKLLLETCSMIKPFVPVTVDGRKWQDHDTEVMANDLRFFNFVPSEKWHGFEGYQEHQYFVDPCKFMLTTPGIDAATGEYTEFGIPATILANFLRENGIIPEKCDLNSILFLLTPAEDMAKLQHLIAQIARFERYVAEDAPLSEVLPNVYRSNEARYRGYTIRQLCQEMHNLYVSYDVKQLQKEMFRQAHFPKAVMNPQDANIEFIRGNVELVPLCKAEGRIAAEGALPYPPGVLCVVPGEVWGGAVQRYFLALEEGINLLPGFSPELQGVYIQEEDDGTKRAYGYVMKQ

**Arginase**

>Consensus1

MKIELRQHKTTRRSSEALTHTHTKRRFALVGAPFNQMGCVTTRANTAAQLRESTDSEPGLSEWMAVRNARWGADIIDAGDVLPSPEVWALLAQLVDGAPDERGFKDQALARYCADLHAMLLTQYRAGRVPLTLGGDHAIAFASVQAALQIYQQEQGKKVAVVWVDAHADCNHTLDSNLHGKPLAMLMNRYPYHGWSVPVERELAAANLFYVGVRDMMPCEYALIRDLGITCFDMALIEQLGFGEIVRRLCRELERDYDHVYLSFDYDALDGSLFRACATPNVGGLSAREALHLVHSVASCAGFIGADFVEYLPERDPDSISKALMVKLMDAVWGYRI

>Consensus2

MKIELRQHKTTSRSSEALTHTHTKLRFALVGAPFNQMGCVTTRANTAAQLRESTDSEPGLSEWMAVRNARWGADIIDAGDVLPSPEVWALLAQLVDGAPDERGFKDQALARYCADLHAMLLTQYRAGRVPLTLGGDHAIAFASVQAALQIYQQEQGKKVAVVWVDAHADCNHTLDSNLHGKPLAMLMNRYPYHGWSVPVERELAAANLFYVGVRDMMPCEYALIRDLGITCFDMALIEQLGFGEVVRRLCRELERDYDHVYLSFDYDALDGSLFRACATPNVGGLSARESLHLVHSVASCAGFIGADFVEYLPERDPDAISKALMVKLMDAVWGYRI

>Consensus3

MKIELRQHKTTRTSSEALTHTHAKRRFALVGAPFNQMGCVTTRANTAAQLRESTDSEPGLSEWMAVRNARWGADIIDAGDVLPSPEVWALLAQLVDGAPDERGFKDQALARYCADLHAMLLTQYRAGRVPLTLGGDHAIAFASVQAALQFYQQEQGKKVAVVWVDAHADCNHTLDSNLHGKPLAMLMNRYPYHGWSVPVERELAAANLFYVGVRDMMPCEYALIRDLGITCFDMALIEQLGFGEVVRRLCRELERDYDHVYLSFDYDALDGSLFRACATPNVGGLSAREALHLVHSVASCAGFIGADFVEYLPERDPDAISKALMVKLIDAVWGYRI

>Consensus9

MKIELRQHKTTRRSSEALTHTHTKRRFALVGAPFNQMGCVTTRANTAAQLRESTDSEPGLSEWMAVRNARWGADIIDAGDVLPSPEVWALLAQLVDGAPDERGFKDQALARYCADLHAMLLTQYRAGRVPLTLGGDHAIAFASVQAALQIYQQEQGKKVAVVWVDAHADCNHTLDSNLHGKPLAMLMNRYPYHGWSVPVERELAAANLFYVGVRDMMPCEYALIRDLGITCFDMALIEQLGFGEIVRRLCRELERDYDHVYLSFDYDALDGSLFRACATPNVGGLSAREALHLVHSVASCAGFIGADFVEYLPERDPDSISKALMVKLMDAVWGYRI

>Consensus10

MKIELRQHKTTSRSSEALTHTHTKLRFALVGAPFNQMGCVTTRANTAAQLRESTDSEPGLSEWMAVRNARWGADIIDAGDVLPSPEVWALLAQLVDGAPDERGFKDQALARYCADLHAMLLTQYRAGRVPLTLGGDHAIAFASVQAALQIYQQEQGKKVAVVWVDAHADCNHTLDSNLHGKPLAMLMNRYPYHGWSVPVERELAAANLFYVGVRDMMPCEYALIRDLGITCFDMALIEQLGFGEVVRRLCRELERDYDHVYLSFDYDALDGSLFRACATPNVGGLSARESLHLVHSVASCAGFIGADFVEYLPERDPDAISKALMVKLMDAVWGYRI

>Consensus11

MKIELRQHKTTRTSSEALTHTHAKRRFALVGAPFNQMGCVTTRANTAAQLRESTDSEPGLSEWMAVRNARWGADIIDAGDVLPSPEVWALLAQLVDGAPDERGFKDQALARYCADLHAMLLTQYRAGRVPLTLGGDHAIAFASVQAALQFYQQEQGKKVAVVWVDAHADCNHTLDSNLHGKPLAMLMNRYPYHGWSVPVERELAAANLFYVGVRDMMPCEYALIRDLGITCFDMALIEQLGFGEVVRRLCRELERDYDHVYLSFDYDALDGSLFRACATPNVGGLSAREALHLVHSVASCAGFIGADFVEYLPERDPDAISKALMVKLIDAVWGYRI

>Consensus12

MKIELRQHKTTRRSSEALTHTHTKRRFALVGAPFNQMGCVTTRANTAAQLRESTDSEPGLSEWMAVRNARWGADIIDAGDVLPSPEVWALLAQLVDGAPDERGFKDQALARYCADLHAMLLTQYRAGRVPLTLGGDHAIAFASVQAALQIYQQEQGKKVAVVWVDAHADCNHTLDSNLHGKPLAMLMNRYPYHGWSVPVERELAAANLFYVGVRDMMPCEYALIRDLGITCFDMALIEQLGFGEIVRRLCRELERDYDHVYLSFDYDALDGSLFRACATPNVGGLSAREALHLVHSVASCAGFIGADFVEYLPERDPDSISKALMVKLMDAVWGYRI

>Consensus13

MKIELRQHKTTSRSSEALTHTHTKLRFALVGAPFNQMGCVTTRANTAAQLRESTDSEPGLSEWMAVRNARWGADIIDAGDVLPSPEVWALLAQLVDGAPDERGFKDQALARYCADLHAMLLTQYRAGRVPLTLGGDHAIAFASVQAALQIYQQEQGKKVAVVWVDAHADCNHTLDSNLHGKPLAMLMNRYPYHGWSVPVERELAAANLFYVGVRDMMPCEYALIRDLGITCFDMALIEQLGFGEVVRRLCRELERDYDHVYLSFDYDALDGSLFRACATPNVGGLSARESLHLVHSVASCAGFIGADFVEYLPERDPDAISKALMVKLMDAVWGYRI

>Consensus14

MKIELRQHKTTRTSSEALTHTHAKRRFALVGAPFNQMGCVTTRANTAAQLRESTDSEPGLSEWMAVRNARWGADIIDAGDVLPSPEVWALLAQLVDGAPDERGFKDQALARYCADLHAMLLTQYRAGRVPLTLGGDHAIAFASVQAALQFYQQEQGKKVAVVWVDAHADCNHTLDSNLHGKPLAMLMNRYPYHGWSVPVERELAAANLFYVGVRDMMPCEYALIRDLGITCFDMALIEQLGFGEVVRRLCRELERDYDHVYLSFDYDALDGSLFRACATPNVGGLSAREALHLVHSVASCAGFIGADFVEYLPERDPDAISKALMVKLIDAVWGYRI

>Consensus15

MKIELRQHKTTRRSSEALTHTHTKRRFALVGAPFNQMGCVTTRANTAAQLRESTDSEPGLSEWMAVRNARWGADIIDAGDVLPSPEVWALLAQLVDGAPDERGFKDQALARYCADLHAMLLTQYRAGRVPLTLGGDHAIAFASVQAALQIYQQEQGKKVAVVWVDAHADCNHTLDSNLHGKPLAMLMNRYPYHGWSVPVERELAAANLFYVGVRDMMPCEYALIRDLGITCFDMALIEQLGFGEIVRRLCRELERDYDHVYLSFDYDALDGSLFRACATPNVGGLSAREALHLVHSVASCAGFIGADFVEYLPERDPDSISKALMVKLMDAVWGYRI

>Consensus16

MKIELRQHKTTSRSSEALTHTHTKLRFALVGAPFNQMGCVTTRANTAAQLRESTDSEPGLSEWMAVRNARWGADIIDAGDVLPSPEVWALLAQLVDGAPDERGFKDQALARYCADLHAMLLTQYRAGRVPLTLGGDHAIAFASVQAALQIYQQEQGKKVAVVWVDAHADCNHTLDSNLHGKPLAMLMNRYPYHGWSVPVERELAAANLFYVGVRDMMPCEYALIRDLGITCFDMALIEQLGFGEVVRRLCRELERDYDHVYLSFDYDALDGSLFRACATPNVGGLSARESLHLVHSVASCAGFIGADFVEYLPERDPDAISKALMVKLMDAVWGYRI

**speE**

>Consensus1

MNAVFARQFTGAKWVGAILASMLVLSSAQAEEKTLHVYNWSDYIAPDTLANFQKASGVKVVYDVFDSNEVLEGKLMAGNTGFDIVVPSASFLERQIKAGVFKPLDKSKLTHYANLDPELLKMLSKHDPDNQYGIPYLWATTGIGYNVDKVKAVLGENAPVDSWDLVLKPENLAKLKSCGVAFLDAPSEVYATVLNYLGKDPNSTDPKDYSGAANDLLLQLRPSVTYFHSSQYINDLANGDVCVAIGWAGDVKQAANRAKEANNGVNVGYSIPKEGALAFFDMLAIPADAKNTDTAYAFMDYLLQPEVIAGVSDAVYYANGNAKATELVKPEIRNDQGIYPTAETRAKMFTLTVKDPKVDRVITRSWTRVKSGQ

>Consensus2

MNAVFARQFTGAKWVGAILASMLVLSSAQAEEKTLHVYNWSDYIAPDTLANFQKASGVKVVYDVFDSNEVLEGKLMAGNTGFDIVVPSASFLERQIKAGVFKPLDKSKLTHYANLDPELLKMLSKHDPDNQYGIPYLWATTGIGYNVDKVKAVLGENAPVDSWDLVLKPENLAKLKSCGVAFLDAPSEVYATVLNYLGKDPNSTDPKDYSGAANDLLLQLRPSVTYFHSSQYINDLANGDVCVAIGWAGDVKQAANRAKEANNGVNVGYSIPKEGALAFFDMLAIPADAKNTDAAYAFMDYLLQPEVIAGVSDAVYYANGNAKATELVKPEIRNDQGIYPTAETRAKMFTLTVKDPKVDRVITRSWTRVKSGQ

>Consensus3

MNAVFARQFTGAKWVGAILASMLVLSSAQAEEKTLHVYNWSDYIAPDTLANFQKASGVKVVYDVFDSNEVLEGKLMAGNTGFDIVVPSASFLERQIKAGVFKPLDKSKLSHYANLDPELLKMLSKHDPDNQYGIPYLWATTGIGYNVDKVKAVLGENAPVDSWDLVLKPENLAKLKSCGVAFLDAPSEVYATVLNYLGKDPNSTDPKDYSGAANDLLLQLRPSVTYFHSSQYINDLANGDVCVAIGWAGDVKQAANRAKEANNGVNVGYSIPKEGALAFFDMLAIPADAKNTDAAYAFMDYLLQPEVIAGVSDAVYYANGNAKATELVKPEIRNDQGIYPTAETRAKMFTLTVKDPKVDRVITRSWTRVKSGQ

>Consensus9

MNAVFARQFTGAKWVGAILASMLVLSSAQAEEKTLHVYNWSDYIAPDTLANFQKASGVKVVYDVFDSNEVLEGKLMAGNTGFDIVVPSASFLERQIKAGVFKPLDKSKLTHYANLDPELLKMLSKHDPDNQYGIPYLWATTGIGYNVDKVKAVLGENAPVDSWDLVLKPENLAKLKSCGVAFLDAPSEVYATVLNYLGKDPNSTDPKDYSGAANDLLLQLRPSVTYFHSSQYINDLANGDVCVAIGWAGDVKQAANRAKEANNGVNVGYSIPKEGALAFFDMLAIPADAKNTDTAYAFMDYLLQPEVIAGVSDAVYYANGNAKATELVKPEIRNDQGIYPTAETRAKMFTLTVKDPKVDRVITRSWTRVKSGQ

>Consensus10

MNAVFARQFTGAKWVGAILASMLVLSSAQAEEKTLHVYNWSDYIAPDTLANFQKASGVKVVYDVFDSNEVLEGKLMAGNTGFDIVVPSASFLERQIKAGVFKPLDKSKLTHYANLDPELLKMLSKHDPDNQYGIPYLWATTGIGYNVDKVKAVLGENAPVDSWDLVLKPENLAKLKSCGVAFLDAPSEVYATVLNYLGKDPNSTDPKDYSGAANDLLLQLRPSVTYFHSSQYINDLANGDVCVAIGWAGDVKQAANRAKEANNGVNVGYSIPKEGALAFFDMLAIPADAKNTDAAYAFMDYLLQPEVIAGVSDAVYYANGNAKATELVKPEIRNDQGIYPTAETRAKMFTLTVKDPKVDRVITRSWTRVKSGQ

>Consensus11

MNAVFARQFTGAKWVGAILASMLVLSSAQAEEKTLHVYNWSDYIAPDTLANFQKASGVKVVYDVFDSNEVLEGKLMAGNTGFDIVVPSASFLERQIKAGVFKPLDKSKLSHYANLDPELLKMLSKHDPDNQYGIPYLWATTGIGYNVDKVKAVLGENAPVDSWDLVLKPENLAKLKSCGVAFLDAPSEVYATVLNYLGKDPNSTDPKDYSGAANDLLLQLRPSVTYFHSSQYINDLANGDVCVAIGWAGDVKQAANRAKEANNGVNVGYSIPKEGALAFFDMLAIPADAKNTDAAYAFMDYLLQPEVIAGVSDAVYYANGNAKATELVKPEIRNDQGIYPTAETRAKMFTLTVKDPKVDRVITRSWTRVKSGQ

>Consensus12

MNAVFARQFTGAKWVGAILASMLVLSSAQAEEKTLHVYNWSDYIAPDTLANFQKASGVKVVYDVFDSNEVLEGKLMAGNTGFDIVVPSASFLERQIKAGVFKPLDKSKLTHYANLDPELLKMLSKHDPDNQYGIPYLWATTGIGYNVDKVKAVLGENAPVDSWDLVLKPENLAKLKSCGVAFLDAPSEVYATVLNYLGKDPNSTDPKDYSGAANDLLLQLRPSVTYFHSSQYINDLANGDVCVAIGWAGDVKQAANRAKEANNGVNVGYSIPKEGALAFFDMLAIPADAKNTDTAYAFMDYLLQPEVIAGVSDAVYYANGNAKATELVKPEIRNDQGIYPTAETRAKMFTLTVKDPKVDRVITRSWTRVKSGQ

>Consensus13

MNAVFARQFTGAKWVGAILASMLVLSSAQAEEKTLHVYNWSDYIAPDTLANFQKASGVKVVYDVFDSNEVLEGKLMAGNTGFDIVVPSASFLERQIKAGVFKPLDKSKLTHYANLDPELLKMLSKHDPDNQYGIPYLWATTGIGYNVDKVKAVLGENAPVDSWDLVLKPENLAKLKSCGVAFLDAPSEVYATVLNYLGKDPNSTDPKDYSGAANDLLLQLRPSVTYFHSSQYINDLANGDVCVAIGWAGDVKQAANRAKEANNGVNVGYSIPKEGALAFFDMLAIPADAKNTDAAYAFMDYLLQPEVIAGVSDAVYYANGNAKATELVKPEIRNDQGIYPTAETRAKMFTLTVKDPKVDRVITRSWTRVKSGQ

>Consensus14

MNAVFARQFTGAKWVGAILASMLVLSSAQAEEKTLHVYNWSDYIAPDTLANFQKASGVKVVYDVFDSNEVLEGKLMAGNTGFDIVVPSASFLERQIKAGVFKPLDKSKLSHYANLDPELLKMLSKHDPDNQYGIPYLWATTGIGYNVDKVKAVLGENAPVDSWDLVLKPENLAKLKSCGVAFLDAPSEVYATVLNYLGKDPNSTDPKDYSGAANDLLLQLRPSVTYFHSSQYINDLANGDVCVAIGWAGDVKQAANRAKEANNGVNVGYSIPKEGALAFFDMLAIPADAKNTDAAYAFMDYLLQPEVIAGVSDAVYYANGNAKATELVKPEIRNDQGIYPTAETRAKMFTLTVKDPKVDRVITRSWTRVKSGQ

>Consensus15

MNAVFARQFTGAKWVGAILASMLVLSSAQAEEKTLHVYNWSDYIAPDTLANFQKASGVKVVYDVFDSNEVLEGKLMAGNTGFDIVVPSASFLERQIKAGVFKPLDKSKLTHYANLDPELLKMLSKHDPDNQYGIPYLWATTGIGYNVDKVKAVLGENAPVDSWDLVLKPENLAKLKSCGVAFLDAPSEVYATVLNYLGKDPNSTDPKDYSGAANDLLLQLRPSVTYFHSSQYINDLANGDVCVAIGWAGDVKQAANRAKEANNGVNVGYSIPKEGALAFFDMLAIPADAKNTDTAYAFMDYLLQPEVIAGVSDAVYYANGNAKATELVKPEIRNDQGIYPTAETRAKMFTLTVKDPKVDRVITRSWTRVKSGQ

>Consensus16

MNAVFARQFTGAKWVGAILASMLVLSSAQAEEKTLHVYNWSDYIAPDTLANFQKASGVKVVYDVFDSNEVLEGKLMAGNTGFDIVVPSASFLERQIKAGVFKPLDKSKLTHYANLDPELLKMLSKHDPDNQYGIPYLWATTGIGYNVDKVKAVLGENAPVDSWDLVLKPENLAKLKSCGVAFLDAPSEVYATVLNYLGKDPNSTDPKDYSGAANDLLLQLRPSVTYFHSSQYINDLANGDVCVAIGWAGDVKQAANRAKEANNGVNVGYSIPKEGALAFFDMLAIPADAKNTDAAYAFMDYLLQPEVIAGVSDAVYYANGNAKATELVKPEIRNDQGIYPTAETRAKMFTLTVKDPKVDRVITRSWTRVKSGQ
